# Differential effects of intra-modal and cross-modal reward value on visual perception: ERP evidence

**DOI:** 10.1101/2021.09.29.462374

**Authors:** Roman Vakhrushev, Felicia Cheng, Annekathrin Schacht, Arezoo Pooresmaeili

## Abstract

Stimuli associated with high reward modulate perception and such value-driven effects have been shown to originate from the modulation of the earliest stages of sensory processing in the brain. In natural environments objects comprise multiple features (imagine a rolling soccer ball, with its black and white patches and the swishing sound made during its motion), where each feature may signal different associations with previously encountered rewards. How perception of such an object is affected by the value associations of its constituent parts is unknown. The present study compares intra- and cross-modal value-driven effects on behavioral and electrophysiological correlates of visual perception. Human participants first learned the reward associations of visual and auditory cues. Subsequently, they performed a visual orientation discrimination task in the presence of previously rewarded visual or auditory cues (intra- and cross-modal cues, respectively) that were concurrently presented with the target stimulus. During the conditioning phase, when reward associations were learned and reward cues were the target of the task, reward value of both modalities enhanced the electrophysiological correlates of sensory processing in visual cortex. During the post-conditioning phase, when reward delivery was halted and previously rewarded stimuli were task-irrelevant, cross-modal value-enhanced behavioral measures of visual sensitivity whereas intra-modal value led to a trend for suppression. A similar pattern of modulations was found in the simultaneously recorded event-related potentials (ERPs) of posterior electrodes. We found an early (90-120 ms) suppression of ERPs evoked by high-value, intra-modal stimuli. Cross-modal cues led to a later value-driven modulation, with an enhancement of response positivity for high-compared to low-value stimuli starting at the N1 window (180-250 ms) and extending to the P3 (300-600 ms) responses of the posterior electrodes. These results indicate that visual cortex is modulated by the reward value of visual as well as auditory cues. Previously rewarded, task-irrelevant cues from the same or different sensory modality have a different effect on visual perception, as intra-modal high-value cues may interfere with the target processing, whereas cross-modal high-value cues boost the perception of the target.

## Introduction

Reward seeking is a fundamental mechanism for survival and one of the main predictors of behavior (Berridge and Kringelbach, 2008; Schultz, 2015). We tend to prioritize what we eat, where we go and what we do based on the expected value of objects or actions learned through experience. A large body of literature has identified a network encompassing the ventral striatum and orbitofrontal cortex to play a key role in learning the associated value of neutral stimuli through experience (Rangel et al., 2008; Schultz, 2000). As natural environments are rich and dynamic, reward information could be conveyed through multiple sensory modalities (e.g., hearing the sound of an approaching ice-cream truck or seeing its characteristic red color both inform us of the possibility of enjoying an ice cream) and reward associations or the goals of the task at hand may change over time (e.g., a red truck can carry other items rather than ice-cream). These features entail a tight interaction between the reward network and the early sensory areas so that stimuli leading to better outcomes such as higher rewards or realization of the goals of the task are prioritized for perceptual processing (Haber, 2011; Pessoa and Engelmann, 2010). In fact, previous research has identified value-driven modulations of neuronal responses in almost all primary sensory areas (Goltstein et al., 2013; Pleger et al., 2008; Rutkowski and Weinberger, 2005; Shuler and Bear, 2006; Stanisor et al., 2013; Weil et al., 2010). In line with the effect of reward on the earliest stages of sensory processing, studies on humans with the use of electroencephalography (EEG) reported reward-related modulations of the visual Event-Related Potentials (ERPs) that are likely to originate from the primary and extrastriate visual cortices (but see also Tankelevitch et al., 2020), including modulations in P1 (80-120 ms) (Bayer et al., 2019; Hammerschmidt et al., 2018; Schacht et al., 2012), N1 (140-220 ms) (Luque et al., 2017) and C1 (∼70 ms) (Bayer et al., 2017; Rossi et al., 2017) ERP componentes. Despite the robust evidence for an influence of reward on early sensory mechanisms, especially in the visual cortex, it has remained unclear how reward information and sensory processing are coordinated across multiple sensory modalities. This is an important question as in natural environments stimuli are typically multisensory and reward information should be hence coordinated across multiple sensory modalities.

Two recent studies tried to bridge this gap and examined cross-modal reward mechanisms, where an auditory stimulus associated with positive reward value affects visual perception (Leo and Noppeney, 2014; Pooresmaeili et al., 2014). During a conditioning phase, participants associated different pure tones with reward values. In a subsequent post-conditioning phase, participants either reported the location (Leo and Noppeney, 2014) or the orientation (Pooresmaeili et al., 2014) of a near-threshold Gabor stimulus in the presence of task-irrelevant auditory tones. Importantly, during the post-conditioning phase, auditory tones did not predict the delivery of reward anymore. The first study (Leo and Noppeney, 2014) showed that participants’ accuracy in determining the location of the near-threshold Gabor was higher in the presence of tones previously associated with a positive reward compared to no reward (neutral). The authors concluded that the rewarded tones enhanced the bottom-up salience of the visual stimuli hence leading to a better detection and more accurate localization of the near-threshold Gabors. Similar results were found in the second study (Pooresmaeili et al., 2014), as it was shown that participants’ orientation discrimination improved in the presence of auditory tones previously associated with high compared to low reward value. With the aid of the simultaneously acquired functional MRI data, it was further shown that the enhanced orientation discrimination is also reflected in the activation patterns of the early visual areas elicited by a specific Gabor orientation (Pooresmaeili et al., 2014). Moreover, in addition to classical reward coding areas (such as Striatum and Orbitofrontal cortex), multisensory regions of the temporal cortex were also modulated by sound values suggesting that they may serve as an intermediary stage to better coordinate the interaction between the primary sensory cortices when high-value stimuli were presented (Pooresmaeili et al., 2014). Subsequent studies (B. A. Anderson, 2016; Kang et al., 2018, 2017; Sanz et al., 2018) used a variety of tasks encompassing visual search or detection or discrimination tasks and confirmed that cross-modal (auditory) reward cues could affect visual perception, albeit the cross-modal reward cues in some cases interfered with the visual task (B. A. Anderson, 2016; Sanz et al., 2018) and in other cases improved it (Kang et al., 2018, 2017). Together, these results provide evidence that reward signals can be broadcasted cross-modally. However, it remains unclear which mechanisms underlie the transmission of reward signals between sensory modalities and their effect on visual perception.

A predominant view posits that reward effects on sensory perception occur through the engagement of attentional mechanisms (Chelazzi et al., 2013; Failing and Theeuwes, 2018; Pessoa, 2015). In this view, rewarding cues receive higher attentional prioritization either through an involuntary, value-driven attentional capture (Anderson et al., 2011a), through voluntary, goal-directed attentional selection (Kiss et al., 2009; Sawaki et al., 2015; Van Den Berg et al., 2014), or through cognitive control mechanisms (Etzel et al., 2016; Pessoa, 2009). In line with this, it has been shown that a reward cue that is aligned with the goal-directed attention in space and in time improves visual performance (Kiss et al., 2009; Kristjánsson et al., 2010), whereas when the rewarded stimulus appears away from the target, it interferes with the target detection or discrimination (Anderson et al., 2013; Della Libera and Chelazzi, 2009). Considering these findings, enhancement of visual perception by co-occurring sounds (Leo and Noppeney, 2014; Pooresmaeili et al., 2014) is unexpected, as auditory tones in these studies were irrelevant to the visual task and could potentially act as a high-reward distractor capturing attention away from the visual target. We consider three possible explanations for the observation that high-reward auditory cues could in some cases enhance visual perception. Firstly, cross-modal auditory reward cues may involve a different type of attentional selection compared to the attentional mechanisms involved in the processing of visual targets (Alais et al., 2006). According to this view, although auditory reward cues capture attention, they do not interfere with the task-relevant visual attention as they are processed through a different attentional mechanism. However, this explanation cannot yet explain why rewarded sounds should *enhance* visual performance. A second possibility is that cross-modal reward cues enhance visual perception by strengthening the audiovisual integration of the auditory and visual components of an audiovisual stimulus. Although audiovisual integration largely occurs automatically (for a review see Calvert and Thesen, 2004), top-down factors such as attention (Alsius et al., 2005; Navarra et al., 2010; Talsma et al., 2007) and recently reward value (Bean et al., 2021; Cheng et al., 2020) have been shown to affect its strength. Through a more efficient integration with the visual cues, auditory reward signals could hence capture attention not only to themselves but also to the whole audiovisual object including the visual target, thereby improving performance. In fact, the boost of integration may be a key characteristic of reward modulation where the association of one sensory property of an object with higher reward spreads to all sensory properties of the same object hence promoting their grouping, as has been shown before (Stanisor et al., 2013). Lastly, cross-modal and intra-modal reward cues could influence visual perception at different stages of sensory processing (Anderson et al., 2011a; Yantis et al., 2012). While intra-modal, task-irrelevant reward cues may interfere with the early bottom-up processing of the visual target, cross-modal reward cues could enhance the later sensory or post-sensory decision stages (Franzen et al., 2020). These possible scenarios can be disentangled by comparing cross-modal reward effects against intra-modal reward effects, i.e., when the reward cue and target cues appear in the same sensory modality. This would also determine whether common or modality-specific mechanisms underlie reward-driven modulation of sensory perception. However, previous studies have not undertaken such a comparison.

The central aim of this study was to provide a better understanding of reward effects across sensory modalities by comparing intra- and cross-modal reward effects, while using the high temporal resolution of the EEG data to delineate different stages of stimulus processing across time. To this end, we employed a task design similar to a previous study (Pooresmaeili et al., 2014), where participants first learned the reward value of visual or auditory cues during a conditioning phase and subsequently performed a visual orientation task in the presence of previously rewarded cues. During the latter post-conditioning phase, auditory and visual reward cues were presented simultaneously with the target stimulus and at the same side of the visual field. Crucially reward cues during the post-conditioning phase were *irrelevant to the task* at hand and *did not predict the delivery of reward* (i.e. no-reward phase). Measuring reward effects at a no-reward phase has proven to be an effective method to separate different modulations of perception, i.e., those related to the long-term associative value of reward cues from those of goal-directed boosts driven by reward-predicting cues or the delivery of the reward itself (Anderson and Yantis, 2013; Failing and Theeuwes, 2015; Maclean and Giesbrecht, 2015; Mine and Saiki, 2015; Rutherford et al., 2010; Watson et al., 2019). Association of task-irrelevant cues, but not the target, with reward values allows dissociating reward effects and effects of selective attention involved in the processing of the target stimulus. Thus, the current study will use task-irrelevant intra- and cross-modal reward cues during a no-reward phase (here referred to as post-conditioning) to measure reward effects across sensory modalities on visual perception.

We hypothesized that cues previously associated with high value should positively influence behavioral performance (i.e., enhancing the target discriminability and reducing the reaction times) and early posterior ERP components (i.e., increasing the amplitudes of P1 and N1 components, and decreasing their latencies). This hypothesis is based on a mechanism where higher reward value not only prioritizes the processing of reward-associated cues but also promotes the integration of different components of an object into a coherent perceptual group. With this, we expected that the reward-related enhancements should spread to other sensory components of the same object, irrespective of the sensory modality of the reward cue. Next, we hypothesized that reward-related modulations of P1 and N1 components in posterior electrodes occur earlier and are stronger when the reward cue is delivered intra-modally (in visual modality) compared to when the reward cue is delivered cross-modally (in auditory modality), as intra-modal effects rely on direct neuronal connections between reward and target sensory representations (Serences, 2008; Serences and Saproo, 2010), whereas cross-modal effects rely on the long-range communication between different brain areas and/or involve intermediate stages (Pooresmaeili et al., 2014). Finally, we hypothesized that in later stages of information processing (i.e. > 250 ms), intra- and cross-modal reward cues elicit similar modulations of ERP amplitudes, both leading to an enhancement of P3 ERP component.

## Material and Methods

### Participants

Thirty-eight participants took part in our experiment (23 women, mean age ± SD: 25.8 ± 5). Two participants were excluded since their performance during the associative reward learning task indicated that they either did not learn the reward associations (N=1) or had difficulties with discriminating the location of the stimuli (N=1). The final sample cconsisted of 36 healthy participants with normal or corrected-to-normal vision who had no history of neurophysiological or psychiatric disorders, according to a self-report. Participants gave written informed consent after the experimental procedures were clearly explained to them. The study was conducted in full accordance with the Declaration of Helsinki and was approved by the local Ethics Committee of the Medical University Göttingen (proposal: 15/7/15).

The sample size, all procedures, and the analysis plan of the study were preregistered (https://osf.io/47wxr/). In brief, the required sample size for this study was calculated based on the effect size observed in a pilot study (N=8). To this end, we averaged the reward effect on the amplitude of the P1 component of the posterior ERPs across cross- and intra-modal cues during post-conditioning. The sample size was then calculated for a within-subjects, paired, two-tailed t-test for a significant difference between high and low value stimuli with alpha = 0.05 and power (1 - Beta) = 0.8. in GPower software (Faul et al., 2007). This power analysis indicated that at least 33 subjects were needed (mean ± SD of effect size: 0.35 ± 0.69 for the difference between high and low value, d_z_ = 0.5097). Based on this and to maintain the counterbalancing of our experimental conditions (4 possible combinations of auditory and visual cues with high or low reward), we aimed for an a priori sample size of N=36 before the data collection started.

### Experimental design

#### Study structure

Each participant completed three phases of the experiment: a pre-conditioning phase (**Figure 1b**), an associative reward learning (conditioning) phase (**Figure 1a**), and finally, a post-conditioning phase (**Figure 1b**): During the pre-conditioning phase, participants performed a visual orientation discrimination task at their discrimination threshold (see Stimuli and equipment) in the presence of the task-irrelevant auditory or visual cues. In the conditioning phase, participants learned reward values of the visual and auditory cues with a cue localization task. Finally, the post-conditioning phase employed the same task as the pre-conditioning but the auditory or visual cues had now been associated with a specific reward value during conditioning. We used the pre-conditioning phase to control for perceptual biases towards visual and auditory cues that were independent of their reward value. Therefore, all effect sizes (Fig. 3-6) are reported after the subtraction of the pre-conditioning data from the corresponding data during post-conditioning.

**Figure 1.**
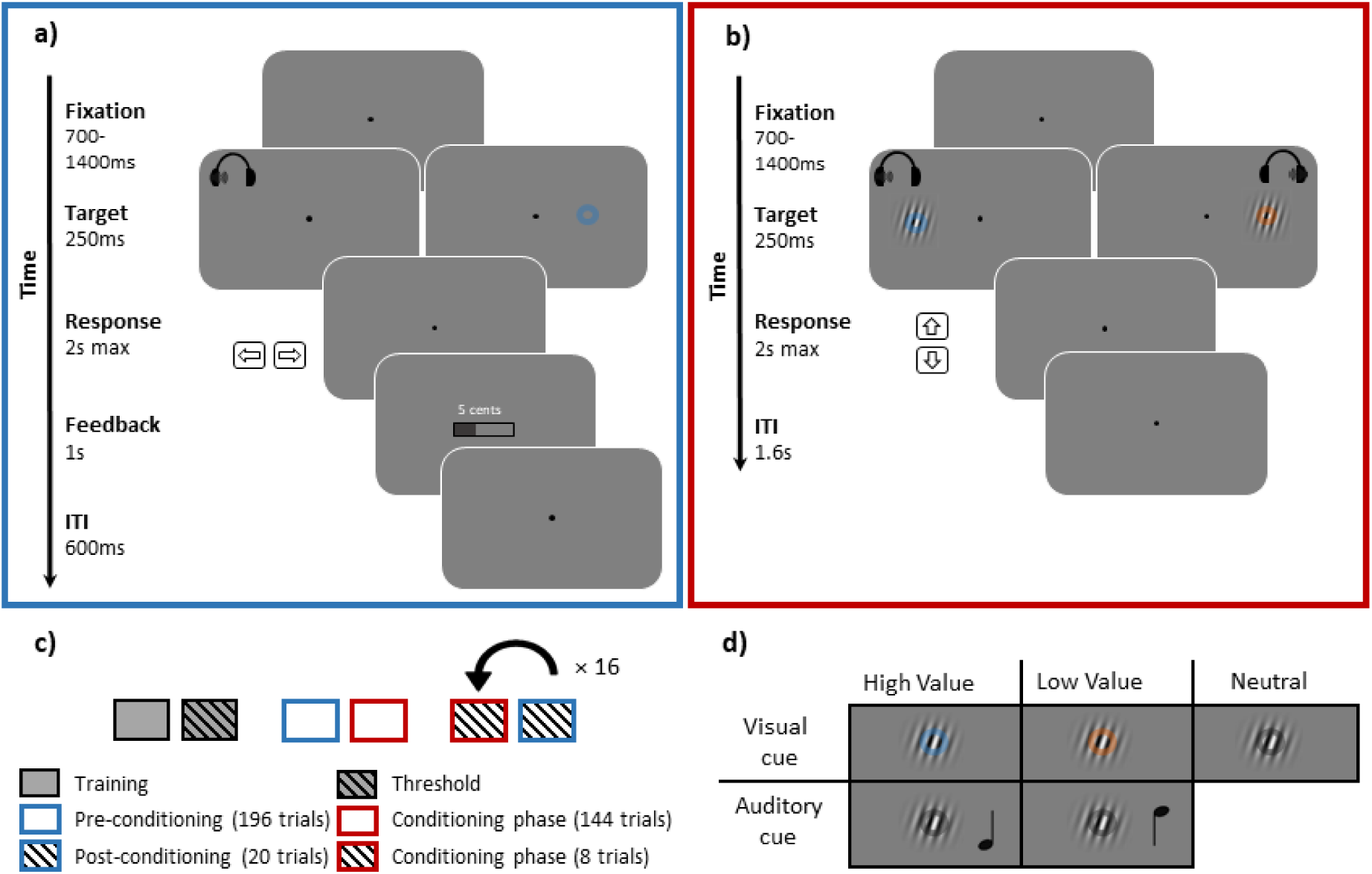
Stimuli and experimental procedure. **a) Reward learning phase (conditioning).** Participants learned the reward association of the visual or auditory cues by performing a localization task and observing their monetary rewards contingent on the color (visual) or pitch (auditory) of the cues. In this task, after an initial fixation period (700-1400 ms), a visual or an auditory cue was presented to the left or right side of the fixation point and participants had to localize them by pressing either left or right arrow buttons of a keyboard (maximum response time 2 s). Here, stimuli in two example trials are shown, one with an auditory cue presented to the left and the other with a visual cue presented to the right. The correct performance led to either a high or a low monetary reward dependent on the identity of the cue (color of visual cues and pitch of auditory cues). **b) Orientation discrimination task employed during the pre- and post-conditioning phases**. To probe the effects of reward value on visual sensitivity, an orientation discrimination task was employed. A trial of this task started with a fixation period (700-1400 ms), followed by the presentation of a peripheral Gabor stimulus (9° eccentricity). Participants were instructed to discriminate the orientation of the Gabor stimulus (clockwise or counterclockwise tilt) by pressing up or down arrow buttons on a keyboard (maximum response duration = 2 s). Concurrent with the Gabor, a visual or auditory cue was also presented (intra- or cross-modal cues, respectively). Intra- and cross-modal cues were irrelevant to the orientation discrimination task and did not predict reward delivery. **c)** The schematic illustration of the different stages of an experimental session. **d)** Design matrix of all stimulus conditions used during the test phase.

#### Conditioning phase

During the conditioning phase (**Figure 1a**), participants reported the location of a peripheral visual (9° eccentricity) or auditory stimulus (sound played in the left or right earphone) by pressing left or right arrow keyboard buttons. Correct responses were followed by a monetary reward, the magnitude of which depended on the respective color or pitch of the visual or auditory cues. Reward information was presented at the screen center both in written format (with digits showing the number of Euro cents obtained) and graphically (a color bar illustrating the ratio of the reward magnitude in current trial to the maximum possible level in the experiment, respectively), and stayed in view for 1.5 s. Rewards were drawn from two Poisson distributions with the mean of 25 cents and SD of 4.8 cents for high-reward trials and a mean of 2 cents and SD 1.4 cents for low-reward trials (minimum and maximum reward was fixed at 35 and 0.4 cents, respectively). Participants were instructed to remember and report the color and the sound pitch that delivered higher rewards at the end of the conditioning phase and at the end of the experiment. The location (left or right), modality (auditory or visual), and identity (color or sound pitch) of the cues were pseudorandomized across trials with the constraint that the same condition could not be repeated more than twice in a row. The association of each cue identity with reward value was counterbalanced across participants.

Participants completed 280 trials with the cue localization task to learn reward associations. From these, 144 trials were presented during an initial longer conditioning block (comprising 36 repetitions of value × modality conditions). Subsequently, 136 trials of the conditioning phase were divided into 16 short blocks (8 trials per block) and then interleaved with short blocks of the post-conditioning phase (20 trials per block) to prevent the extinction of the reward associations (**Figure 1c**, also see under *Pre- and post-conditioning phases*).

#### Pre- and Post-conditioning phases

During pre- and post-conditioning phases, participants performed an orientation discrimination task (**Figure 1b**). Each trial started with a fixation period (700-1400 ms) followed by a Gabor stimulus. The Gabor stimulus was displayed at 9° eccentricity on the left or right side from the fixation point for 250 ms (see also Stimuli and equipment). Simultaneously with the Gabor stimulus, an ipsilateral visual (intra-modal) or auditory (cross-modal) cue appeared. Participants were instructed to focus on judging the orientation of the Gabor and ignore irrelevant visual and auditory cues. During these phases, correct responses in the orientation discrimination task granted them a fixed reward shown at the end of the experiment. These instructions aimed to emphasize that previously rewarded cues were not relevant to the task and did not lead to differential immediate rewards anymore.

In total, participants performed 160 trials in the pre-conditioning phase and 320 trials in the post-conditioning phase, with 32 and 64 repetitions of each of the value (high, low, neutral) × modality (intra-modal, cross-modal). In the neutral condition, the Gabor was presented without an auditory tone or color and served as a control condition to characterize the behavioral performance and evoked potentials, irrespective of the reward value (**Figure 1c-d**).

#### Stimuli and equipment

Visual reward cues consisted of a transparent ring overlaid on the Gabor patch (0.44° in diameter, 0.17 pixels thick, alpha 50%, color in HSV = [20, 0.4, V] for orange, [250, 0.4, V] for blue and [0, 0, V] for grey colors and V was calibrated for each participant to produce isoluminance, intra-modal conditions contained either a blue or an orange ring whereas cross-modal and neutral conditions contained a grey ring). Auditory reward cues were pure monaural tones, lateralized to the left or right earphone (pitch = 350 Hz or 1050 Hz and sound level pressure = 70 dB, adjusted with a digital sound-level meter; SL-100, Voltcraft, Hirschau; separately for each tone and delivered through in-ear earphones ER-3C, Etymotic Research Inc.). Auditory cues were not localized in space as we aimed to keep their characteristics identical to a previous study (Pooresmaeili et al., 2014). Gabor stimuli were Gaussian-windowed sinusoidal gratings with an SD of 0.33°, a spatial frequency of 3 cycles per degree, 2° diameter, and 50% contrast, displayed on a gray background of 34.4 cd/m^2^ luminance. Before the experiment, participants performed a color adjustment procedure to match the brightness of the visual reward cues. After that, they underwent a short training session to get familiar with the orientation discrimination task (Number of trials = 36, **Figure 1c**). After training, participants’ orientation discrimination threshold for a performance of 70% was determined using a QUEST procedure, followed by the main experiment (**Figure 1c**). Throughout the experiment, trials were repeated in case participants made an eye movement (> 2° displacement of gaze from the fixation dot during fixation or target period), used an undesignated button to respond, or did not respond at all to maintain equal number of repetitions across conditions.

Data collection was done in a darkened, sound-attenuated, and electromagnetically shielded chamber. Participants sat 91 cm away from a 22.5-inch calibrated monitor (ViewPixx/EEG inc., resolution = 1440 × 980 pixels, refresh rate = 120 Hz) with their heads resting on a chinrest. Stimulus presentation was controlled in Psychophysics toolbox-3 (Brainard, 1997) in MATLAB (version R2015b) environment. Eye movements were recorded with an Eyelink 1000 eye tracker system (SR Research, Ontario, Canada) in a desktop mount configuration, recording the right eye at a sampling rate of 1000 Hz. Electrophysiological data were recorded from 64 electrodes (BrainVision Recorder 1.23.0001 Brain Products GmbH, Gilching, Germany; actiCap, Brain Products GmbH, Gilching, Germany), online referenced to TP9, digitized at 1000 Hz, and amplified with a gain of 10,000. Electrode impedances were kept below 10kΩ.

### Data Analysis

#### Analysis of the behavioral data

For behavioral assessment of visual sensitivity during the orientation discrimination task, we used participants’ d-prime scores (d’). D-prime was measured based on the probability of hits and false-alarms, as d’ = Z(P_Hit_) - Z(P_FA_), where one of the tilt directions was arbitrarily treated as “target-present” as in formal Signal Detection Theory analysis of discrimination tasks (Macmillan and Creelman, 1991; Leo et al., 2011). Extreme values of P_Hit_ or P_FA_ were slightly up- or down-adjusted (i.e., a probability equal to 0 or 1 was adjusted by adding or subtracting 0.001, respectively). Reaction times in all phases of the experiment were calculated as the elapsed time between the onset of the target stimulus and the participant’s response. The resulting response times were averaged for each experimental condition (including correct and incorrect responses).

Outliers were removed from the behavioral and ERP data of each participant (0.38% ± 0.6 SD across subjects). A trial was considered to be an outlier if the gaze fixation during the target presentation period was suboptimal (eye position >0.9° from the fixation point) or the reaction time was faster than 10 ms.

#### Analysis of Event-Related Potentials (ERPs)

The EEG data was imported and processed offline using EEGLAB (Delorme and Makeig, 2004), an open-source toolbox running under the MATLAB environment. First, an automatic bad channel detection and removal algorithm was applied (using EEGLAB’s pop_rejchan method; threshold = 5, method = kurtosis). Later, data of each participant was band-pass filtered with 0.1 Hz as the high-pass cutoff and 40 Hz as the low-pass cutoff frequencies. After this, the epochs were extracted by using a stimulus-locked window of 1900 ms (−700 to 1200 ms) and subjected to an Independent Component Analysis (ICA) algorithm (Delorme & Makeig, 2004). Blinks and eye-movement artifacts were automatically identified and corrected using an ICA-based automatic method, implemented in the ADJUST plugin of EEGLAB (Mognon et al., 2011). Bad channels were interpolated using the default spherical interpolation method of the EEGLAB toolbox. Next, data were re-referenced to the average reference. To calculate ERPs, epochs were extracted using a window of 900 ms (−100 to 800 ms) time-locked to the onset of the target, and baseline-corrected using the pre-stimulus time interval (−100 to 0 ms).

Our ERP analysis focused mainly on the latencies and amplitudes of the visual P1 and N1 components of the visual and auditory reward cues averaged across a region of interest (ROI) comprising four posterior electrodes (PO7/PO8, O1/O2). As auditory and visual cues have different response latencies and reward associations may also affect response latencies, the following method was used to tailor P1 and N1 amplitude measures to each specific condition. ERP components were defined individually for each condition as a 30 ms window around the peak centered on the grand average ERP wave across participants. The peak of P1 component was defined as the most positive peak within 70-170 ms window, and the peak of N1 component was defined as the most negative peak within 180-250 ms window. Latencies of P1 and N1 peaks were measured by finding the corresponding positive or negative peaks of the ERPs within the corresponding time windows for each participant. Finally, modulations of the P3 component were quantified as the average ERP amplitude between 300-600 ms across all electrodes of the midline ROI (Fz, FCz, Cz, CPz, and Pz).

To determine the onset of intra- and cross-modal reward effects, we calculated the difference between high- and low-value ERPs of each cue type within a 10 ms moving window against their baseline difference (i.e. within 100 ms before the stimulus onset). Amplitude differences between 50-800 ms after stimulus onset that reached significance (uncorrected p < 0.05) were considered as significant value-driven modulations. To account for the inflated probability of type I errors that appear from application of multiple comparisons, only results that were significant in two or more consecutive time windows will be reported (Talsma and Woldorff, 2005).

All procedures described above strictly followed our pre-registered analysis plan. However, after data acquisition and visual inspection of the ERPs, we undertook the following modifications and exploratory analyses. Firstly, in the post-conditioning phase, we observed ERP modulations in time intervals that preceded our pre-registered time windows. To examine these early effects, we performed an exploratory analysis in a time window between 90-120 ms after the stimulus onset (referred to as the PA component). The timing of this component corresponds to the timing of the earliest positive peak of visual ERPs observed in previous studies (Luck et al., 1994; Mangun, 1995). It should be noted that in our experiments, reward cues and the Gabor stimulus were presented at the same time. As such, whereas the highest positive peak of P1 that has been the basis of our estimates of P1 component is likely to reflect the responses to both reward cues and the Gabor, the earlier PA component may reflect primarily the response to the reward cues. This conclusion is also supported by inspecting the PA responses to the neutral stimulus depicted in Figure 4a and Supplementary Figure 4a, where PA responses start to emerge only after the reward associative learning (in post-conditioning phase) when participants pay attention to the identity of different cues. The second exploratory analysis was to inspect the stimulus-evoked contra-lateral ERPs. As visual areas are known to have a higher sensitivity to the contralateral stimuli (Woldorff et al., 1997), we also tested the responses of the posterior ROI to contralateral stimuli (i.e., responses of O1 and PO7 to the stimuli on the right visual hemifield and O2 and PO8 to the stimuli on the left visual hemifield) and measured the correlation of contralateral responses with behavior (Figure 6, Supplementary Figure 4), as exploratory analyses. Finally, ERP modulations occurring at a time window overlapping with the P3 responses (300-600 ms) planned to only be analyzed in midline electrodes were also inspected in the posterior ROI (O1, O2, PO7, PO8) during both conditioning and post-conditioning phases.

#### Statistical Analysis

We used 2 by 2 repeated measures ANOVAs (rm-ANOVAs) to test the effect of *Reward Value* (high, low) and *Modality* (visual, auditory) on behavioral (d’ and RT) and electrophysiological responses of the conditioning and post-conditioning phases. To remove the effect of perceptual biases that participants may have for different colors or tone pitches prior to the learning of cues’ reward associations, we corrected the behavioral and ERP results of each participant by subtracting the data of each condition in pre-conditioning from the data of post-conditioning. The corrected results were then subjected to rmANOVAs. Planned, paired t-tests were used for pairwise comparisons. Effect sizes in rmANOVAs are reported as partial eta-squared 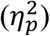 and in pairwise comparisons as Cohen’s d; i.e. d_z_ (Lakens, 2013).

## Results

### Conditioning phase

#### Behavioral results

Participants’ performance in this task was near perfect as accuracies for both cues were > 95% (99% ± 1% for auditory and 100%±0% visual cues). Analysis of reaction times (RTs) revealed a significant main effect of modality (F_(1,35)_ = 70.44, P < 0.001, η_p_^2^ = 0.67). Participants localized visual cues (mean ± s.e.m.: 501 ± 12 ms) faster than auditory cues (mean ± s.e.m.: 584 ± 16 ms), in line with the superior performance of vision in localization tasks (Bertelson and Radeau, 1981; Pick et al., 1969). We did not observe a main effect of reward value (F_(1,35)_ < 1); however, an interaction was found between reward value and modality (F_(1,35)_ = 7.68, p = 0.009, η_p_^2^ = 0.18). For auditory cues, high reward value slowed down responses (mean ± s.e.m.: 589 ± 17 ms and 580 ± 16 ms for high and low value cues, respectively), while in the visual modality, high-value cues sped up responses (mean ± s.e.m.: 496 ± 13 ms and 506 ± 12 ms for high- and low-value cues, respectively). The effect of reward value separately in each modality did not reach significance (t(35) = -2.02, p = 0.051, d_z_ = 0.34 and t(35) = 1.52, p= 0.137, dz = 0.253 for visual and auditory cues, respectively).

#### ERP responses: P1, N1, P3

We next examined whether the posterior ROI (PO7/PO8 and O1/O2, reflecting the activity of occipital and parietal visual areas) that we had planned to use for probing reward effects during the main orientation discrimination task exhibits reliable reward modulations (ERP amplitude and or latency) during reward associative learning (**Figure 2**). The rationale for examining visual ERPs in the conditioning phase is twofold. Firstly, in this phase reward modulations in the posterior ROI serve as an index of how value information affects sensory processing when reward cues are the target of the task (task-relevant). Secondly, of particular interest are reward modulations evoked by auditory cues, as such modulations would indicate that reward information is broadcasted to visual cortex even in the absence of concomitant visual stimulation.

**Figure 2.**
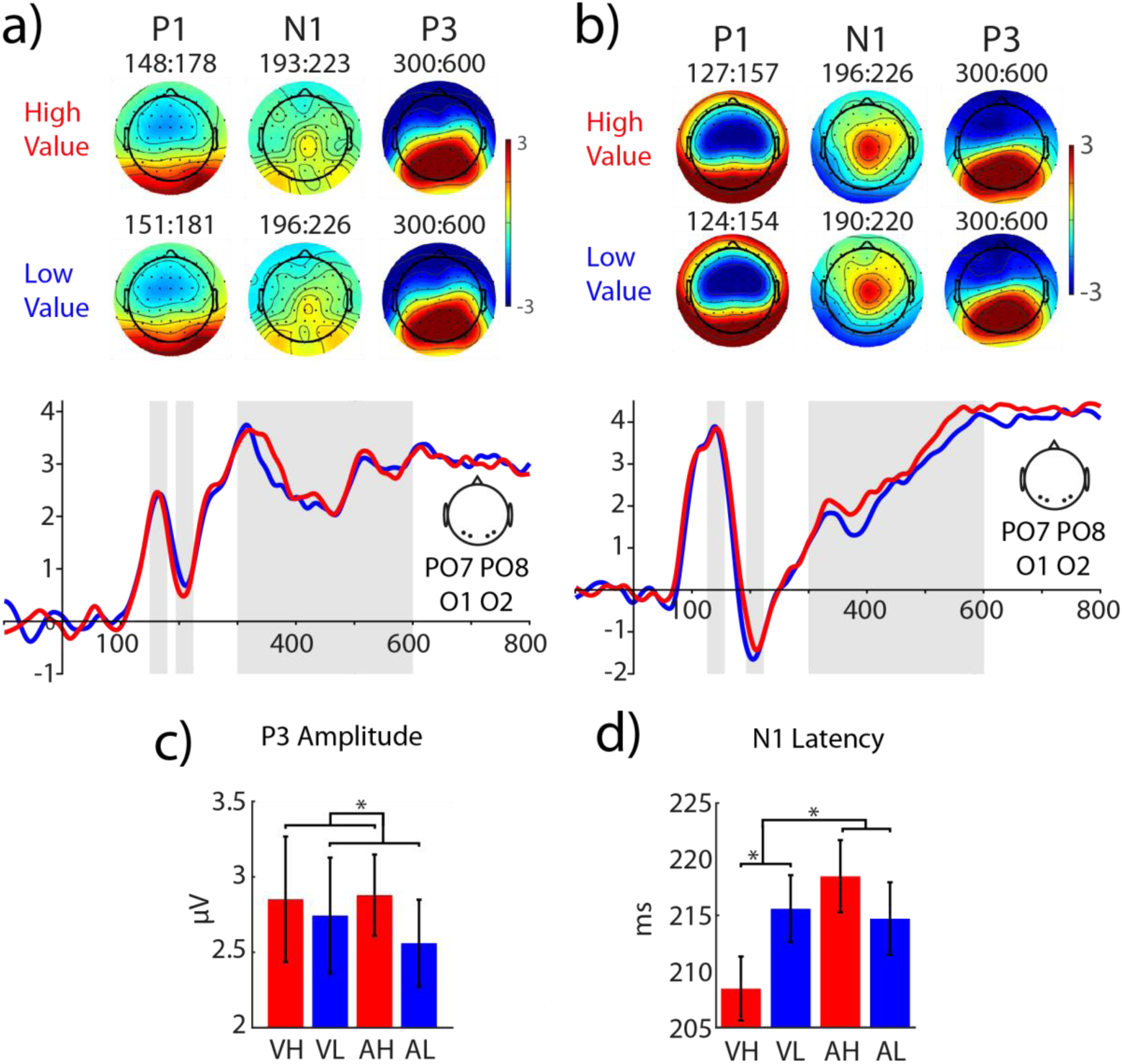
ERP responses of the posterior ROI to the visual and auditory reward cues during the conditioning phase. **a)** ERPs of visual reward cues with high (red traces) and low (blue traces) values measured in a posterior ROI (O1, O2, PO7 and PO8). The shaded grey areas correspond to 30 ms windows around the peak of P1 (70-170 ms) and N1 (180-250 ms) used to estimate the amplitude of these components for each condition (here the window is averaged across high and low value conditions) and the window used to estimate P3 responses (300-600 ms). The topographic distribution of P1, N1 and P3 components are shown for each reward value condition. **b)** Same as **a** for auditory reward cues. **c-d**) analysis of the amplitude and latency of the ERP components revealed significant value-driven modulations for the amplitude of P3 shown in **c** and the latency of the N1 component shown in **d**. Visual High Value: VH; visual Low Value: VL; Auditory High Value: AH; Auditory Low Value: AL.

According to the rmANOVA, ***P1 amplitudes*** were impacted by modality (F_(1,35)_ = 12.89, p < 0.001, η_p_^2^ = 0.27), with higher amplitudes for auditory (mean ± s.e.m: 3.66 ± 0.28 µV) than visual cues (mean ± s.e.m: 2.27 ± 0.24 µV). Effects of reward values (F_(1,35)_ < 1, *p* = 0.919) and the modality by reward interaction (F_(1,35)_ < 1, *p* = 0.971) did not reach significance. Analysis of ***P1 latency*** revealed a main effect of modality (F_(1,35)_ = 53.1, *p* < 0.001, η_p_^2^ = 0.60), reflecting an earlier onset of P1 responses for auditory (mean ± s.e.m.: 130.7 ± 2 ms) compared to visual cues (mean ± s.e.m.: 155.3 ± 2.7 ms). Neither a main effect of reward value (F_(1,35)_ = 1.19, *p* = 0.284) nor an interaction between reward value and modality (F_(1,35)_ = 2.72, *p* = 0.108) was found.

Analysis of ***N1 amplitudes*** in the conditioning phase revealed a main effect of modality (F_(1,35)_ = 17.42, p < 0.001, η_p_^2^ = 0.33), again with larger amplitudes for auditory (mean ± s.e.m: -1.36±0.35 µV) compared to visual cues (mean ± s.e.m: 0.73±0.32 µV). Again, neither the main effect of reward value nor the reward by modality interaction reached significance (all ps > 0.1). ***N1 latency*** was not modulated by modality (F_(1,35)_ = 1.92, *p* = 0.175) or by reward value (F_(1,35)_ < 1, *p* = 0.477). However, a significant interaction was found between reward value and modality (F_(1,35)_ = 7.07, p = 0.012, η_p_^2^ = 0.17), reflecting different direction of value-driven modulation of N1 latencies in the two modalities. Whereas visual high value cues significantly sped up the N1 peak (mean ± s.e.m.: 208.47 ± 2.86 ms and 215.58 ± 2.99 ms and for high and low value cues respectively, t(35) = -2.54, *p* = 0.016, d_z_ =0.424), auditory high-value cues tended to slow down N1 responses (mean ± s.e.m.: 218 ± 3.3 ms and 214.7 ± 3.2 ms for high and low value cues respectively, t(35) = 1.12, p = 0.27, Cohen’s d =0.19).

Analysis of ***P3 amplitude*** in the posterior ROI (PO7, O1, O2, PO8), quantified between 300 and 600 ms, revealed a main effect of reward value (F_(1,35)_ = 4.82, *p* = 0.035, η_p_^2^ = 0.121), reflecting larger amplitudes for high-reward cues (Mean ± s.e.m.: 2.86 ± 0.29 µV) than for low-reward cues (Mean ± s.e.m.: 2.65 ± 0.3 µV). The main effect of modality (F_(1,35)_ < 1, *p* = 0.816) and an interaction effect between modality and value (F_(1,35)_ < 1, *p*= 0.458) were non-significant.

We note that this analysis is exploratory as in our pre-registered plan we intended to only examine the P3 component in midline electrodes (Pz, CPz, Cz, FCz, Fz) in a time window of 300-600 ms. However, examination of the topographic distribution of these ERP modulations within this window suggested a rather posterior topography than the expected more centrally distributed P300 (Figure 2a-b). Hence, we decided to also examine P3 amplitudes in a posterior ROI (O1, O2, PO7, PO8). Similar motivational modulation of posterior P3-like responses has been observed in previous studies (Baines et al., 2011).

In summary examination of the posterior ROI (O1, O2, PO7, and PO8) revealed a significant interaction of reward value and reward modality on the latency of N1 responses, with a significant speeding up of ERP responses to high-value compared to low-value visual cues, and opposite trend for auditory N1 responses. In the P3 window, a main effect of reward value was found across all high-compared to low-reward value conditions. Taken together, these results indicate that the higher associative value of both visual and auditory cues enhances their sensory processing in visual areas during the associative learning phase of our experiment.

### Post-Conditioning phase

#### Behavioral results

To assess the behavioral visual sensitivity, participants’ d-prime scores (d’ Post *minus* d’ Pre) were subjected to an ANOVA with reward value (high or low) and modality (intra- or cross-modal) as independent factors (Table 2 and **Figure 3**). This analysis revealed no main effect of modality (F_(1,35)_ < 1, *p* = 0.99) or reward value (F_(1,35)_ < 1, *p* = 0.35). Importantly, an interaction was found between modality and reward value (F_(1,35)_ = 5.75, p = .022, η_p_^2^ = 0.14). Whereas cross-modal reward value enhanced visual sensitivity (mean ± s.e.m: 0.44 ± 0.19, for the difference between high and low value stimuli, corrected for pre-conditioning, t(35)=2.31, p = 0.027, d_z_ = 0.38), intra-modal reward value led to a decrement in visual sensitivity (mean ± s.e.m.: -0.18 ± 0.19, for the difference between high and low value stimuli, corrected for pre-conditioning). The performance decrement for high-compared to low-value intra-modal condition however, did not reach statistical significance (t(35)=-0.97, p = 0.34, d_z_ = 0.162). Analysis of the reaction times (RT) revealed a trend for an overall slowing down of responses for high-compared to low-value stimuli, but this effect did not reach statistical significance (F_(1,35)_ = 2.36, *p* = 0.133). Other main and interaction effects were non-significant (all Fs < 1, all ps > 0.1).

**Table 1.**
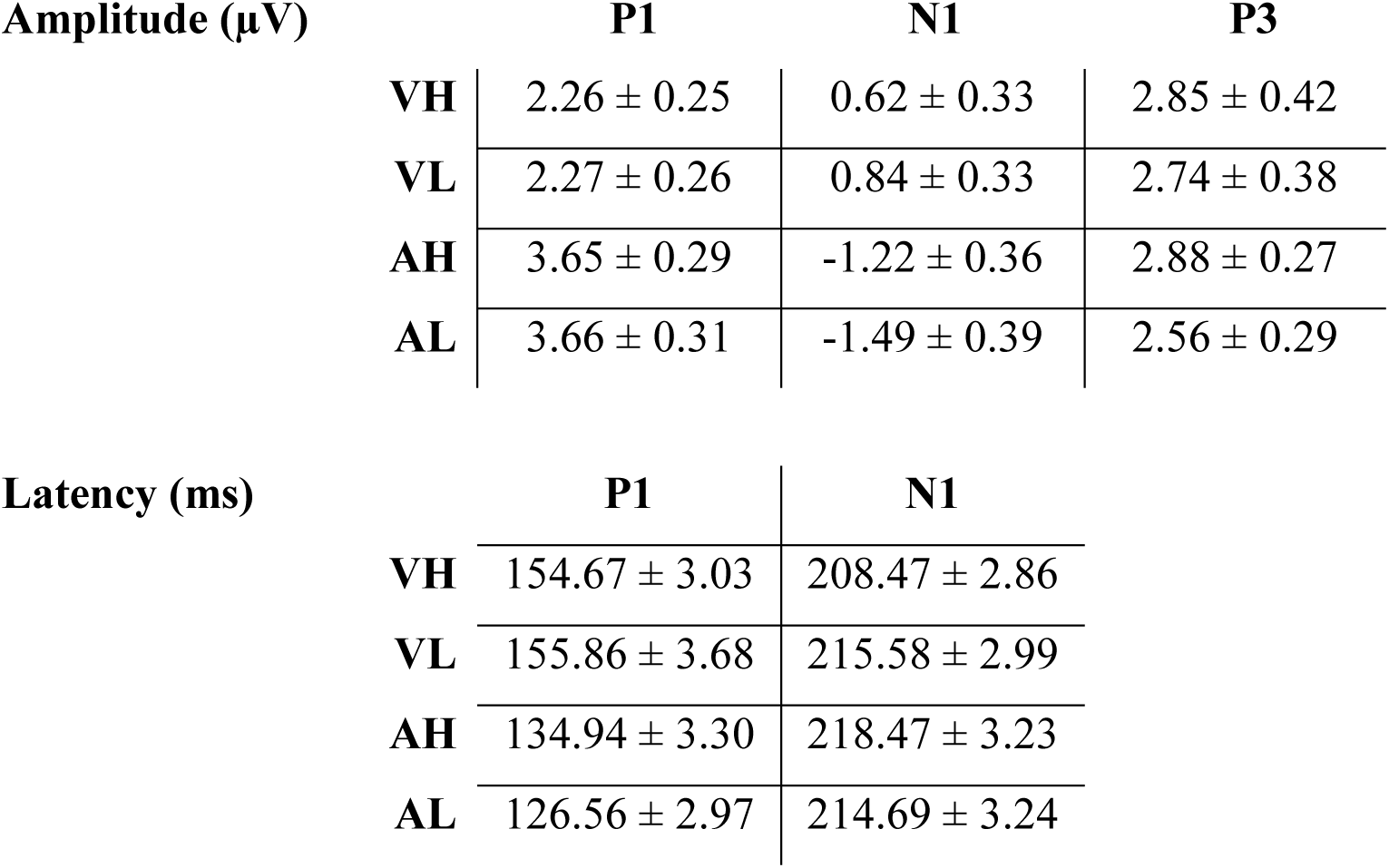
Amplitude and latencies of visual ERP components evoked by visual high value (VH), visual low value (VL), auditory high value (AH), and auditory low value (LA) cues during the conditioning phase, measured in the posterior ROI.

**Table 2.** Summary of behavioral and ERP data during the pre- and post-conditioning phases (the gray and white cells, respectively) for intra-modal (IH and IL, high and low values), cross-modal (CH and CI, high and low value), and neutral (Neut) conditions. Significant pairwise comparisons are shown in bold fonts (p < 0.05).

|  | *Pre-conditioning* |  |  |  |  | *Post-conditioning* |  |  |  |  |
| --- | --- | --- | --- | --- | --- | --- | --- | --- | --- | --- |
|  | IH | IL | CH | CL | Neut | IH | IL | CH | CL | Neut |
| RT (ms) | 950 ±25.3 | 968 ±28.2 | 955 ±28 | 961 ±27.8 | 955 ±29.7 | 923 ±21.7 | 917 ±23.4 | 930 ±22.6 | 922 ±20.9 | 906 ±22.8 |
| d' | 2.18 ±0.17 | 2.18 ±0.18 | 1.94 ±0.16 | 2.24 ±0.21 | 2.05 ±0.16 | 1.89 ±0.13 | 2.06 ±0.14 | **1.96 ±0.16** | **1.82 ±0.14** | 1.81 ±0.14 |
| PA (μV) | 0.62 ±0.29 | 0.16 ±0.26 | 3.52 ±0.40 | 3.89 ±0.41 | 0.09 ±0.24 | **0.09 ±0.15** | **0.39 ±0.17** | 3.36 ±0.34 | 3.30 ±0.36 | 0.40 ±0.17 |
| P1 (μV) | 2.28 ±0.36 | 2.35 ±0.35 | 5.56 ±0.43 | 6.03 ±0.39 | 1.98 ±0.34 | 1.84 ±0.26 | 2.07 ±0.30 | 4.81 ±0.34 | 5.00 ±0.36 | 1.93 ±0.26 |
| N1 (μV) | 0.93 ±0.41 | 0.99 ±0.39 | -1.29 ±0.46 | -0.67 ±0.48 | 0.42 ±0.42 | 0.38 ±0.35 | 0.74 ±0.35 | **-0.70 ±0.40** | **-0.94 ±0.41** | 0.91 ±0.36 |
| P3 (μV) | 2.65 ±0.41 | 2.70 ±0.43 | 3.44 ±0.47 | 4.23 ±0.49 | 1.98 ±0.46 | 2.58 ±0.46 | 2.48 ±0.45 | **3.91 ±0.47** | **3.76 ±0.49** | 2.52 ±0.45 |
| P1 (ms) | 151 ±3.63 | 152 ±3.78 | 141 ±3.08 | 145 ±2.78 | 147 ±4.15 | 147 ±4.01 | 147 ±4.33 | 140 ±3.62 | 136 ±3.50 | 151 ±3.26 |
| N1 (ms) | 214 ±3.85 | 213 ±3.71 | 214 ±2.82 | 213 ±2.81 | 211 ±3.59 | 209 ±3.29 | 215 ±3.34 | 213 ±2.66 | 210 ±2.33 | 212 ±3.65 |

**Figure 3.**
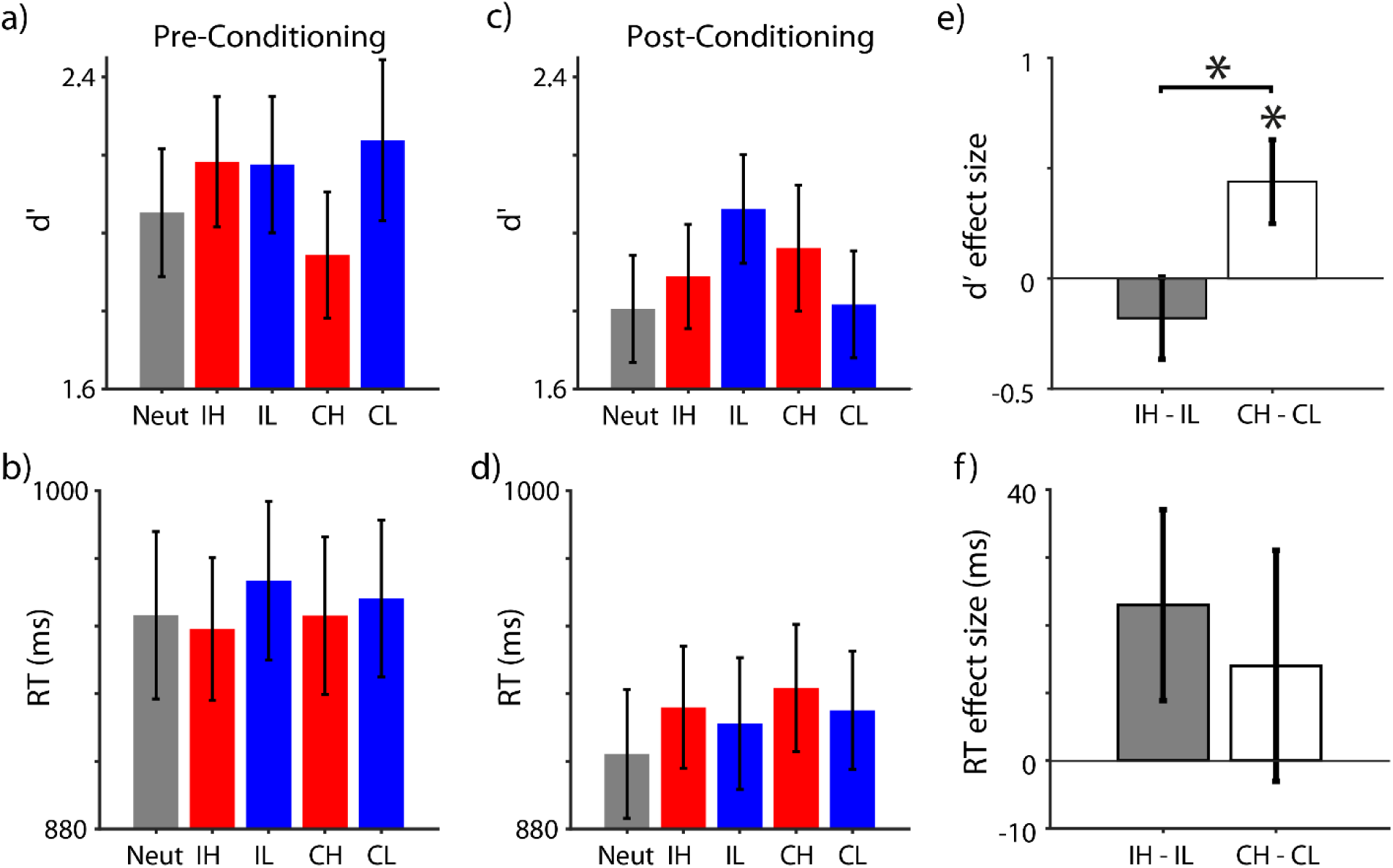
Behavioral performance in the orientation discrimination task. **a)** Visual sensitivity (d-prime: d’) during the pre-conditioning phase for different conditions (Neutral: Neut; Intra-modal High Value: IH; Intra-Modal Low Value: IL; Cross-modal High Value: CH; and Cross-Modal Low Value: CL). **b)** Same as **a**, for reaction times (RT). **c)** Visual sensitivity during the post-conditioning phase, i.e. after the associative reward value of different cues were learned. **d)** Same as **c** for RTs. **e)** Effect size for d’ modulations, measured as the difference in d′ between high- and low-value conditions corrected for their difference during pre-conditioning, in intra- and cross-modal cue types (grey and white bars, respectively). **f)** Same as **e** for reaction times (RT). * marks the significant effects (p < 0.05). Error bars are s.e.m.

In summary, during the post-conditioning phase, cross-modal high-value cues improved the visual sensitivity of orientation discrimination while intra-modal, high-value cues showed a trend for interference compared to low-value cues. Reaction times were not significantly affected by the reward value of either type.

#### ERP responses in posterior ROI

We next examined the responses of the posterior ROI to the Gabor stimuli when it appeared together with task-irrelevant visual or auditory cues during the orientation discrimination task (**Figure 4, Figure 5**, and **Table 2**). Examination of ***P1 amplitudes*** with a two-way rmANOVA revealed a main effect of modality (F_(1,35)_ = 5.65, p =0.023, η_p_^2^ = 0.14), corresponding to larger P1 amplitudes in the cross-modal condition than in the intra-modal condition (see **Table 2**). The main effect of reward value (F_(1,35)_ < 1, *p* = 0.831) and the interaction between reward value and modality did not reach significance (F_(1,35)_ < 1, *p* = 0.501). Analysis of ***P1 latencies*** did not reveal any effect of modality (F_(1,35)_ < 1, *p* = 0.935), value (F_(1,35)_ < 1, *p* = 0.342), or their interaction (F_(1,35)_ = 1.2, *p* = 0. 287).

**Figure 4.**
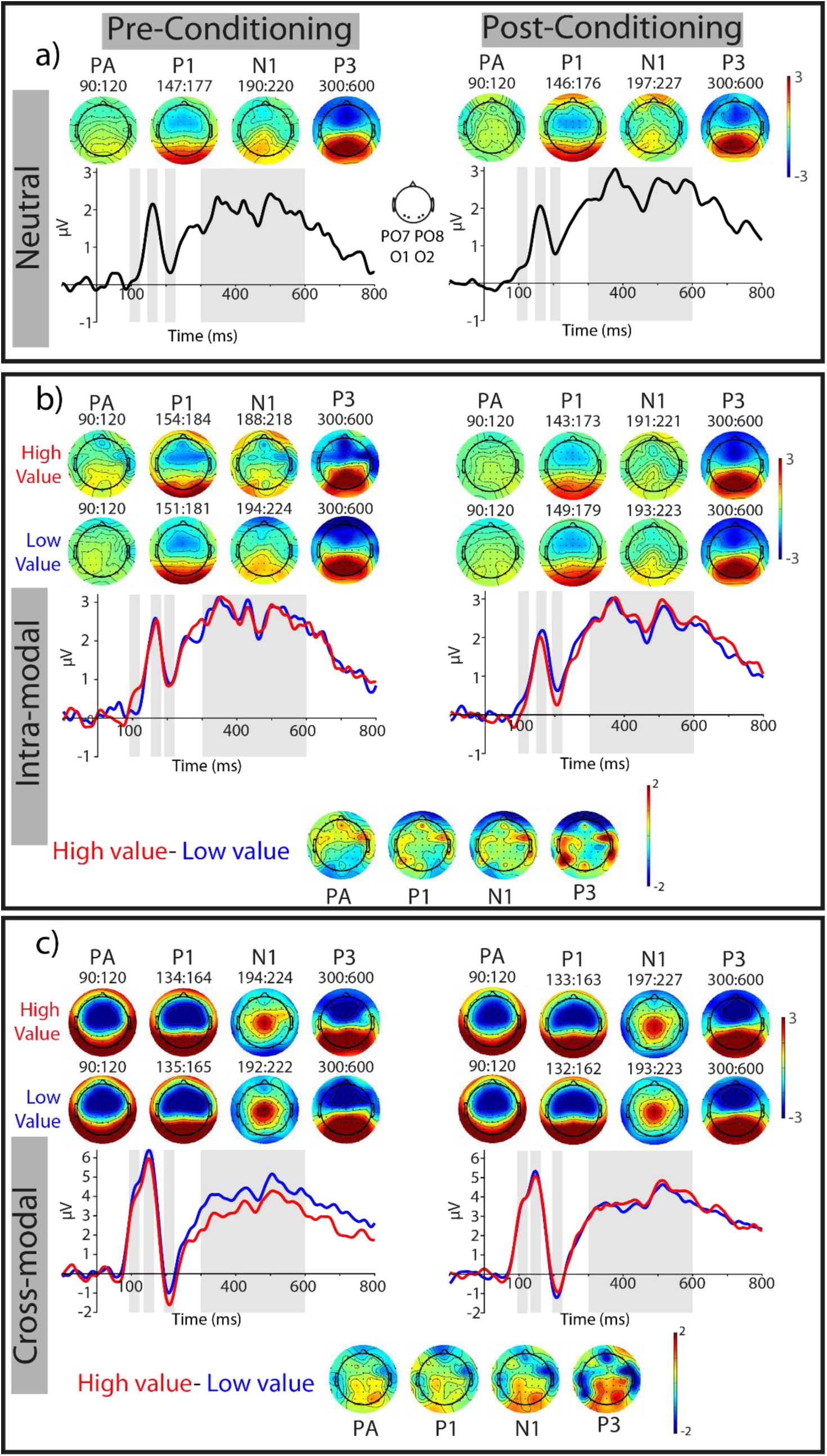
ERP responses of the posterior ROI (PO7, PO8, O1, O2) during the visual orientation discrimination task. **a)** ERPs elicited by the neutral condition during pre- and post-conditioning (illustrated on the left and right, respectively). The topographic distributions of PA (90-120 ms), P1 (most positive peak 70-170 ms), N1 (most negative peak 180-250 ms, and P300 (300-600 ms) components, measured in respective grey shaded areas of the ERP time courses are also shown. **b)** ERPs in the presence of task-irrelevant, intra-modal reward cues, with high (red traces) and low (blue traces) values. The corresponding topographic distribution of each component is shown for each reward value condition. To test the significance of value-driven modulations, the difference between high- and low-value conditions was corrected for their pre-conditioning difference. The topographic distributions of these corrected modulations are shown for each component (the lowermost topographic maps). See also Figure 5c. **c)** Same as **b** for ERPs in the presence of task-irrelevant, cross-modal reward cues.

**Figure 5.**
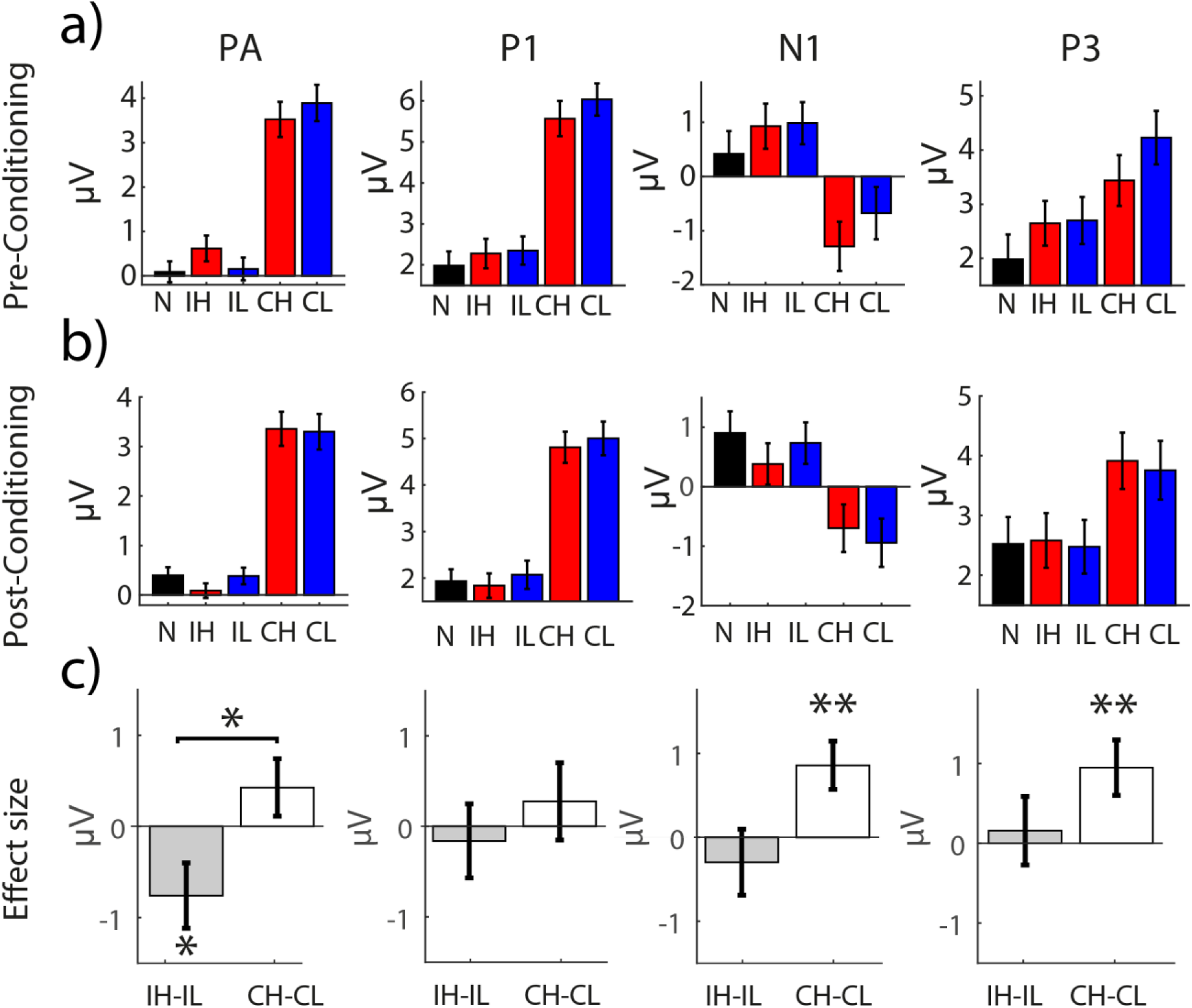
Mean amplitude of different ERP components of the posterior ROI (PO7, PO8, O1, O2) during the visual orientation discrimination task. **a)** mean amplitude of PA, P1, N1 and P3 responses evoked by neutral (N), intra-modal high value (IH), intra-modal low value (IL), cross-modal high value (CH) and cross-modal low value (CL) conditions, during pre-condoning. **b)** Same as **a**, for ERPs during post-condoning. **c)** Corrected effect sizes, measured as the difference between high- and low-value conditions corrected for their difference during pre-conditioning, for intra- and cross-modal cue types (grey and white bars, respectively). * mark significant differences (p< 0.05) and ** mark significant differences (p< 0.01). Error bars are s.e.m.

**Figure 6.**
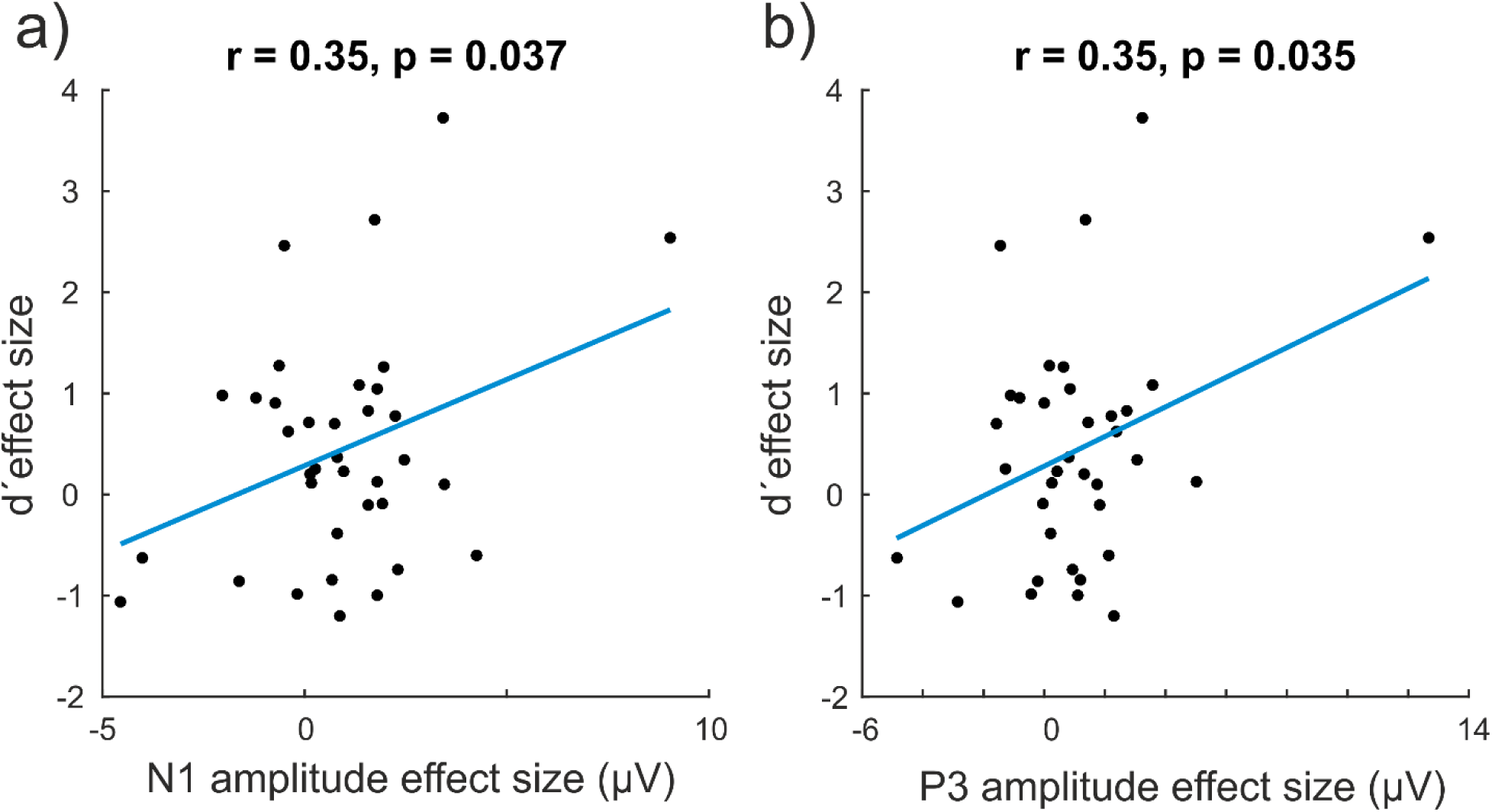
Correlation between electrophysiological and behavioral effects of reward value. **a)** Correlation between reward modulation of contralateral N1 amplitude and d’ in cross-modal condition. Effect size corresponds to the difference of high- and low-value conditions corrected for their difference during pre-conditioning. **b)** Same as **a** for the contralateral P3 amplitude.

Analysis of the ***N1 amplitude*** revealed a trend for an effect of modality (F_(1,35)_ = 3.6, p = 0.066, η_p_^2^ = 0.093), with intra-modal reward cues evoking less negative N1 peak than cross-modal reward cues (see **Table 2**). The main effect of reward value did not reach significance (F_(1,35)_ = 2.04, p = 0.162) but a trend was found for an interaction between reward value and modality (F_(1,35)_ = 4.09, p = 0.051, η_p_^2^ = 0.105), where high value cross-modal cues decreased N1 negativity compared to intra-modal condition (see **Table 2** and **Figure 4-5**). Planned, pairwise comparisons showed that cross-modal high value condition significantly decreased (t(35) = 2.98, p= 0.005, d_z_ = 0.5, **Figure 5c**) the N1 negativity compared to the low value condition. The difference between intra-modal, high and low value conditions did not reach statistical significance (p= 0.45). **Latency analysis of N1** component did not reveal any effects of modality (F_(1,35)_ < 1, *p* = 0.935), value (F_(1,35)_ < 1, *p* = 0.363) or their interaction (F_(1,35)_ = 2.17, *p* = 0.150).

Analysis of the P3 component in midline electrodes (300-600 ms, corrected for pre-conditioning) revealed no main effect of modality, reward value, electrode, or their interaction (all ps > 0.1). Although post-hoc inspection of the ERP responses of midline electrodes (**Supplementary Figures 2** and **3**) suggests that value-driven modulations occur in other time windows than our pre-registered intervals, we decided to adhere to our a priori plan and do not explore these modulations any further.

Based on our pre-registered analysis plan, we next examined the earliest and latest time points, where the reward modulations of each cue type were significant using a moving window analysis (see Methods). This analysis revealed an earlier onset of value-driven modulations for intra-modal compared to cross-modal conditions in the posterior ROI. High- and low-value ERP responses of the intra-modal conditions differed significantly within the P1 window, i.e. between 101 and 114 ms. The cross-modal value-modulations started later, at 167 ms, and remained significant until 223 ms, thus overlapping with the N1 time window (**Figure 4**). Based on these results and visual inspection of the ERPs, we performed an exploratory analysis on the amplitude of evoked responses in a time window between 90-120 ms, where early value-driven modulation was found for intra-modal conditions. This amplitude modulation occurred earlier than our pre-registered measure of P1 component and will be referred to as the PA component throughout. A two-way ANOVA on PA amplitudes revealed a significant value × modality interaction effect (F_(1,35)_ = 5.12, p = 0.03, η_p_^2^ = 0.128), but no main effect of reward value or modality (both Fs < 1 and ps > 0.1). Post-hoc pairwise comparisons (**Figure 5c**) revealed a significant suppression of PA amplitude for high-compared to low-value intra-modal condition (t(35) =-2.1, p = 0.04, d_z_ = 0.35). The value-driven modulation in the cross-modal conditions was in opposite direction but did not reach statistical significance (t(35) = 1.35, p = 0.185, d_z_ = 0.22). We wondered whether the negative modulation of PA responses by intra-modal reward value reflects a suppression of high-reward stimuli or rather an enhancement of the low-reward conditions. To this end, we contrasted the PA amplitudes of intra-modal high- and low-value conditions, respectively, against the neutral condition that had not undergone reward associative learning (see also **Figure 4-5**). These comparisons revealed a significant suppression of PA responses evoked by intra-modal, high-value compared to neutral stimuli (t(35) = -2.07, p = 0.044, d_z_ = 0.35). The intra-modal, low-value stimuli, however, were not significantly different from the neutral condition (t(35) = 0.25, p = 0.801, d_z_ = 0.042). These results suggest that intra-modal high-value stimuli undergo active suppression compared to both intra-modal low value and neutral conditions occurring very early after the onset of stimuli.

We next performed an exploratory analysis on the ERPs of the posterior ROI in a later time window (300-600 ms) corresponding to the ***P3 component***. This analysis revealed a main effect of reward value (F_(1,35)_ = 5.63, *p* = 0.023, η_p_^2^ = 0.138). The main effect of modality and the modality by reward interaction were not significant (both ps > 0.1). Planned, pairwise comparisons revealed a significant enhancement of P3 responses by cross-modal high-compared to low-value condition (t (35) = 2.72, P=0.01, d=0.45). In the intra-modal condition, P3 amplitudes did not differ between the two reward conditions (t(35) = 0.37, *p* = 0.72, d_z_ = 0.061). Together, examination of late ERPs in the posterior ROI mainly indicated a value-driven response enhancement for cross-modal condition (**Figure 5c**).

In our pre-registered analysis plan, we had intended to compare the ERP effects of the correct and error trials. However, after the data acquisition, we noticed that we could not have a noise-free estimation of the ERPs of error trials, as in some cases errors were too infrequent (Mean ± SD: 6.6 ± 2.4 and 13.6 ± 5.4, for number of error trials in pre- and post-conditioning, respectively). Therefore, we did not undertake this pre-registered analysis, albeit we tested whether we obtain similar results when we only include correct trials in our analysis (for which we had sufficient number of trials; i.e.,> 25 trials for each condition and each phase). Our analysis showed that overall the pattern of results remains unchanged when only correct trials were included in our analyses and in most cases effect sizes were larger than when all trials were included in our analysis (Supplementary Text and **Supplementary Figure 5**).

Finally, due to the retinotopic organization of visual cortex, responses to lateralized stimuli are strongest in contralateral cortical regions; therefore, we tested potential value-driven modulations of ERPs of each condition in contra-lateral posterior ROIs separately (**Supplementary Figure 4**). Results from these analyses were highly similar to those, where ERPs in the posterior ROI were averaged across both hemispheres (see also the Supplementary Information).

Taken together, the ERP effects provide evidence for an interaction effect between intra- and cross-modal reward cues, with intra-modal cues leading to an early suppression and cross-modal reward cues producing a later response enhancement for high-compared to low-value stimuli.

#### Correlation of behavioral and ERP effects

We next measured the correlation between value-driven amplitude modulations of components for which a significant reward effect was found (i.e., PA, N1 and P3, see **Figure 5**) and changes in behavioral indices (d’ and RT). This analysis did not reveal any significant correlation (all ps> 0.1). We wondered whether the lack of a correlation with behavior, particularly with d’ is due to averaging the responses of contralateral and ipsilateral electrodes in our ROI. Due to the strong retinotopic organization of early visual areas, the reward modulations of lateralized stimulus are strongest and most reliably correlated with behavior on the contralateral visual areas (Pooresmaeili et al., 2014). To this end, we performed an exploratory analysis where the responses of the contralateral electrodes of the posterior ROI measured in each time window (corresponding to PA, N1, and P3) were correlated with the behavioral d’ of the corresponding condition. This analysis revealed a significant positive correlation between value-driven modulations of the contralateral ERPs in N1 (r = 0.347, p = 0.037, Pearson Correlation, d_z_ = 0.74) and P3 (r = 0.352, p = 0.035, Pearson Correlation, d_z_ = 0.75) windows and behavioral d-primes of the cross-modal stimuli. These results hence indicate that the cross-modal reward effects on behavior may indeed rely on the value-driven response modulations of the contralateral visual areas. However, we note that these results should be treated with caution, as they reflect results obtained from an exploratory analysis and were not corrected for multiple comparisons.

## Discussion

In the current study we tested whether intra-modal and cross-modal reward cues exert similar effects on the processing of visual stimuli. To this end, we examined behavioral and electrophysiological responses to reward cues during a conditioning phase when reward associations were learned and during a post-conditioning phase when rewards were not delivered anymore. Importantly, during post-conditioning reward cues were irrelevant to the visual discrimination task that participants had to perform. In the conditioning phase, we found that intra- and cross-modal reward cues affect latency of the N1 responses over the posterior electrodes differently: while intra-modal reward cues significantly sped up N1 response, there was a trend for deceleration of N1 responses by cross-modal reward cues. However, in the P3 component, we found higher amplitudes for both intra- and cross-modal high value cues. In the post-conditioning phase, similarly to conditioning phase P3-like responses of the posterior ROI were enhanced for high value cues of both types. Importantly, we found much earlier reward effects within 90-120 ms (P1 window), where intra-modal high value reward cues suppressed ERP responses compared to the low value reward cues. The reward effects from cross-modal cues, on the other hand, led to a later modulation at a time point corresponding to the N1 component. Behavioral results were in line with electrophysiological findings; we found an interaction between the reward value and the modality of the reward cue: cross-modal reward cues significantly enhanced visual sensitivity whereas intra-modal reward cues showed a trend for suppression. Interaction effects of reward value with modality within P1 and N1 windows suggest that intra-modal and cross-modal reward values exert different effects on visual perception under settings employed in our study (i.e., task-irrelevant reward cues during a no-reward phase). Similar reward effects of intra- and cross-modal conditions in the P3 window indicate that beyond the differential effect of modality on early visual responses, later reward effects are independent of sensory modality. In the following sections, we will discuss possible mechanisms that might explain such interaction effects at the early ERP components.

### Reward-driven modulations during conditioning

In this phase, a visual target predictive of higher rewards sped up reaction times and early cue-evoked neural responses, particularly in the N1 window. This result is similar to early modulations of visual ERPs (i.e. <250 ms) observed in previous studies (Baines et al., 2011; Hammerschmidt et al., 2018; Hickey et al., 2010; Luque et al., 2017, 2015; Maclean and Giesbrecht, 2015). Auditory stimuli also evoked strong responses in the visual cortex even in the absence of concomitant visual stimulation (Feng et al., 2014; McDonald et al., 2013; Weinberger, 2007; Wikman et al., 2019). However, reward signals from auditory stimuli did not significantly affect the amplitude or latency of early visual ERP components when auditory stimuli were presented alone, albeit a trend was found for deceleration of N1 responses. Lack of cross-modal reward effects during the conditioning phase may indicate that reward information does not transfer automatically when there is no incoming visual information. A trend for deceleration of N1 responses and reaction times by cross-modal reward cues may also be due to the type of the task employed during conditioning (i.e., a localization task) as auditory modality is generally less sensitive to spatial properties of the stimuli than vision (Bertelson and Radeau, 1981; Pick et al., 1969).

The similarity of intra- and cross-modal reward effects on later P3 component indicates that at the later stages of sensory processing reward information integrates into a coherent reward representation across various sources irrespective of the unique characteristics of those sources. On the other hand, during the early stages of sensory processing, privileged processing of rewarded stimuli only prioritizes cues that are most suited for the task at hand, i.e., vision in case of a task requiring localization. These results are overall explainable within the framework of reinforcement learning, where association of a stimulus with reward results in value-driven modulations at 2 time points (Luque et al., 2015): an early modulation of neural responses for stimuli associated with higher value within the first 200-250 ms after the stimulus onset (Luque et al., 2015; Maclean and Giesbrecht, 2015); and a later reward modulation primarily the P3 component related to anticipation of the reward delivery (Goldstein et al., 2006; Luque et al., 2017, 2015; Pornpattananangkul and Nusslock, 2015; Yeung and Sanfey, 2004).

### Reward-driven modulations during post-conditioning

The results obtained in the post-conditioning phase showed a differential pattern of reward-related modulation for cross-modal and intra-modal cues. Specifically, we observed an improvement of behavioral measures of visual sensitivity for high compared to low-value cross-modal cues. These results are in line with the reported facilitatory effects of cross-modal value on visual processing observed in human psychophysical and neuroimaging studies (Leo et al., 2011; Pooresmaeili et al., 2014). As a first attempt to characterize the electrophysiological correlates of this effect, our study demonstrated that the behavioral advantage conferred by high-reward auditory cues is accompanied by a modulation of visual ERPs. As these modulations were corrected for differences potentially occurring due to physical stimulus features (i.e. tone pitches) already during the pre-conditioning phase, they most probably reflect effects driven by associative learning of the reward values.

Cross-modal reward modulations in our study occurred later than attentional effects observed in some of the previous studies of cross-modal attention (Störmer et al., 2009), where response modulations were found in P1 window. However, in these experiments auditory tones were presented prior to the visual target and could therefore modulate the early ERP responses. In fact, when auditory and visual components of an audiovisual object were presented together, attentional effects occurred at a later time window around 220 ms (Busse et al., 2005; Zimmer et al., 2010). Whereas reward-driven boost of attention may to some extent account for cross-modal reward effects in our study, the direction of response modulations suggests that additional mechanisms may also be involved. Specifically, we found a reduction in N1 negativity after learning of the reward associations, which is different from an enhanced P1 positivity and N1 negativity that has been observed in studies of cross-modal attention (Busse et al., 2005; Störmer et al., 2009; Zimmer et al., 2010). One possible mechanism is a reward-driven enhancement of audiovisual integration, which is in line with recent findings demonstrating a role of reward in multisensory integration (Bean et al., 2021; Cheng et al., 2020). This mechanism can also explain the direction of ERP modulations, as previous studies found that an audiovisual stimulus elicits response modulations of visual ERPs mainly in N185 window, with a reduction of the negativity of N185 component compared to the unimodal stimuli (Giard and Peronnet, 1999), which is similar to the pattern of modulations we observed for cross-modal rewards. The reduction in N1 negativity may indicate that audiovisual integration enhances the gain of the neural responses of visual cortex, hence reducing the energy (i.e. N1 amplitude) needed to process the same load of sensory information, an example of sub-additive cross-modal interactions reported before (Molholm et al., 2002; Teder-Sälejärvi et al., 2005, 2002). An enhanced integration between auditory reward cues and visual target in our task potentially reduces the distracting effect of task-irrelevant sounds on visual discrimination while promoting the spread of privileged processing from rewarded sounds to the visual target. Later modulations by cross-modal compared to intra-modal rewards suggests that a putative reward-driven enchantment of audiovisual integration may first occur in multimodal areas such as in Superior Temporal Sulcus (STS), being then fed back to visual cortex, a proposal in line with the findings of neuroimaging studies of cross-modal value and emotion effects on vision (Pooresmaeili et al., 2014; Watson et al., 2014). In addition to the above mechanisms (reward-driven boost of attention and audiovisual integration), the later value-driven modulations in P3 window may indicate that the cross-modal reward effects also rely on post-sensory and decisional stages (Franzen et al., 2020).

The effects of intra-modal reward cues on visual perception were contrary to our a-priori hypothesis that intra-modal and cross-modal reward stimuli should have similar facilitatory effects on visual perception. This hypothesis was based on previous findings (Stanisor et al., 2013) that reported a spread of reward enhancement effects from one component of an object to its other parts (here from colored circles signaling reward to the Gabor target), akin to the spread of object-based attention (Duncan, 1984; Pooresmaeili and Roelfsema, 2014). Contrary to our prediction, we found that intra-modal reward stimuli interfered with sensory processing of the visual target, which was reflected in suppressed ERP responses at an early time window of 90-120 ms elicited by high rewards compared to both low reward and neutral conditions. The interference effect that we observed is in line with the findings of several studies where value-driven effects during visual search were investigated (Anderson, 2013; Anderson et al., 2011b, 2011a; Chang and Egeth, 2021; Feldmann-Wüstefeld and Schubö, 2013; Gong et al., 2016; Hickey et al., 2009; Rusz et al., 2020, 2018; Yantis et al., 2012). These experiments have consistently reported that presentation of a high reward stimulus at the target location speeds up visual search and enhances target-evoked responses, whereas presentation of high reward stimuli at a the distractor location captures attention away from the target and interferes with the search task. Given these previous findings, we superimposed reward cues on the target to boost the processing of all object elements at that location. However, since the visual discrimination task was performed on a different feature of the object (i.e. orientation) than the defining feature of the reward cue (color), high reward visual cues may have captured attention away from the target feature, hence interfering with the target processing. A similar interference effect has been observed in studies where a certain feature of an object was predictive of its reward but this feature was incongruent with the goal of the task, e.g. in many tasks employing the Stroop Effect (Bustamante et al., 2021; Grégoire and Anderson, 2019; Krebs et al., 2013, 2011, 2010).

In light of these previous studies, we propose four possible mechanisms for the observed intra-modal, reward-related suppression. Firstly, it is possible that as a result of an enhanced response to the high-reward distractors (Itthipuripat et al., 2019) some form of local inhibition is exerted on the adjacent stimuli, thereby decreasing the overall responses to target and rewarding distractors that were at the same location. Such a center-surround inhibition around an attended feature has been observed in studies of feature-based and spatial attention (Hopf et al., 2006; Mounts, 2000; Störmer and Alvarez, 2014). Secondly, it is possible that the suppression is a reflection of the higher processing load of high-reward cues (Engelmann, 2009) and the capacity limitation of attentional processing. This mechanism could therefore result from a mixture of enhanced processing of distractors in some trials and decrement of processing of *target+distractor* in other trials due to the depletion of attentional resources. Thirdly, it is possible that the suppression is due to cognitive control mechanisms that actively inhibit the processing of intra-modal reward cues with a resultant spillover of inhibition to the target as also observed in previous studies of feature-based attention (Nobre et al., 2006). The fourth possibility is that reduction of neural responses in visual cortex is a general characteristic of reward-related modulations occurring due to direct dopaminergic inputs to visual cortex (Arsenault et al., 2013) or value-driven attentional gain control mechanisms (Shapcott et al., 2020) that adjust signal-to-noise ratio of neural responses. These scenarios may all contribute to the suppression of responses to intra-modal high reward cues that we observed, although the early onset of the suppressive effects best matches the first two scenarios. Future studies will be needed to tease apart these possibilities.

In summary, our study showed that intra- and cross-modal reward stimuli affect visual perception during a no-reward (post-conditioning) phase differently. In line with our a priori hypothesis intra-modal reward cues affected visual sensory processing early, thus the modulation took place within the visual cortex. Cross-modal reward effects, on the other hand, occurred much later, and exhibited characteristics of audiovisual integration, thus being likely to rely on feedback from multimodal areas, such as STS. We did not expect to find a difference between the direction of modulations evoked by intra- and cross-modal reward stimuli. However, ERP analysis of P1 and N1 components revealed the difference between these conditions, most importantly an early suppression of intra-modal high reward condition. We suggested a number of factors that could contribute to the latter effect: a local suppression induced by reward-irrelevant cues on adjacent target stimuli, a competition for processing resources between target and reward stimuli, active inhibition due to cognitive control mechanisms, or a general mechanism of reward modulation where dopamine as a result of high-reward stimuli adjust signal-to-noise ratio, thus suppress the neuronal response. The first two mechanisms are most favorable, but subsequent studies are needed to tease them apart. Overall, our study provides evidence for a value-driven plasticity in visual perception that can be induced not only by signals from the visual modality but also cross-modally. Embedding cross-modal rewards in visual tasks is a promising tool to assist vision, especially in the face of visual impairments, through boosting cross-modal advantages conferred by another intact sensory modality.

## Acknowledgments

We thank Adem Saglam for his help with programming of the experiment, Franziska Ehbrecht for her help with the data collection, and Jessica Emily Antono for her valuable comments on the manuscript. This work was supported by an ERC Starting Grant (no: 716846) to AP.

## Authors’ contributions

RV and AP conceptualized and designed the experiment. All authors participated in the pre-registration of the study. RV conducted the experiments. RV, FC, and AP analyzed the data. RV and AP interpreted the results and wrote the first draft of the manuscript. All authors revised the manuscript.

## Supplementary Information to

### Supplementary Text

#### ERP results when contralateral posterior ERPs were examined

We evaluated the amplitude of ERP components (PA, P1, N1, and P3) in contralateral electrodes (**Supplementary Figure 4**). To this end, responses of O2 and PO8 to the stimuli presented on the left hemi-field and responses of O1 and PO7 to the stimuli on the right hemi-field were measured. The contralateral ERPs were then averaged across the two sides. Two-way ANOVAs with reward value and modality as independent factors and amplitude of each component as dependent factor revealed only a trend for an interaction between reward value and modality in N1 window (F_(1,35)_ = 3.19, p = 0.083, η_p_^2^ = 0.083), corresponding to a decrease in N1 negativity for cross-modal, high-compared to low-value condition (mean ± s.e.m: 0.41 ± 0.28 and -0.49 ± 0.39 for high- and low-value stimuli respectively, t(35) = 2.3, p = 0.025, d_z_ = 0.389). Furthermore, a main effect of reward value was found in P3 window (F_(1,35)_ = 9.14, p = 0.005, η_p_^2^ = 0.207). Planed pairwise comparisons revealed a significant enhancement of contralateral P3 responses for high-compared to low-value cross-modal cues (mean ± s.e.m: 0.5 ± 0.28 and -0.59 ± 0.44, for high-compared to low-value cross-modal cues respectively, t(35) = 2.388, p = 0.022, d_z_ = 0.398).

#### P3 responses of midline electrodes during conditioning

Previous studies have reported robust reward modulations of P3 responses of midline electrodes (Pz, CPz, Cz, FCz and Fz) time-locked to the presentation of reward cues, both for both visual and auditory cues (Gehring and Willoughby, 2002; Glazer et al., 2018; Krugliakova et al., 2019, 2018; Van Den Berg et al., 2014). Based on these and our pre-registered plan, we next inspected the ERP responses of the midline electrodes (**Supplementary Figure 1**), time-locked to the presentation of the reward cues in the P3 window (300-600 ms). The frontocentral electrodes are most sensitive to auditory signals (Fz, FCz), whereas the posterior midline electrodes are most sensitive to visual cues (Cz, Pz) or both (Cz), based on the studies reported above. Therefore, we expected to find a significant difference in P3 amplitudes for stimuli of different modalities, reflecting the different topography of areas processing visual or auditory information. In addition, we also expected an effect of reward value for one or both modalities across these electrodes. We did not find an overall robust value-driven modulation based on our preregistered analysis plan. Specifically, a rmANOVA revealed a main effect of modality (F_(1,35)_ = 4.64, P = 0.038, η_p_^2^ = 0.117), a main effect of electrode (F_(4,140)_ = 124.73, p < 0.001, η_p_^2^ = 0.781) and a trend for modality and electrode interaction (F_(4,140)_ = 3.53, p = 0.054, η_p_^2^ = 0.092), as expected. Other comparisons did not reach significance: main effect of value (F_(1,35)_ = 1.6, p = 0.214), interaction between value and modality (F_(1,35)_ = 3.19, p = 0.083), interaction between value and electrode F(1,35) < 1, p = 0.381) and interaction between factors value, modality and electrode (F_(1,35)_ = 1.34, p = 0.054). Although the inspection of ERP responses (**Supplementary Figure 1**) suggests value-driven modulations in some of these electrodes for visual (esp. Pz), auditory (esp. Fz), or both cues types (Cz), in order to adhere to our preregistered plan, we did not further investigate these effects.

#### ERP results when only correct trials were included

We examined the ERP responses of the posterior ROI when only correct trials of each condition were included in our analysis (**Supplementary Figure 5**). To this end, we performed rmANOVAs on the amplitude of PA, P1, N1 and P3 and the latency of P1 and N1 components of the ERPs with reward value (high or low) and modality (intra-modal or cross-modal) as factors. Overall, we obtained the same results as results obtained with all trials, albeit some of our reported were even stronger when only correct trials were included.

In PA (90-120 ms) window we found an interaction effect between factors modality and reward value (F(1,35) = 10.54, p = 0.003, η_p_^2^ = 0.231). Intra-modal high-value cues suppressed PA amplitudes compared to low-value cues (mean ± s.e.m: -0.58 ± 0.35 and 0.41 ± 0.27, for the PA amplitude of high-compared to low-value cues, t(35) = -2.475, p = 0.018, d_z_ = 0.412). Cross-modal high value cues on the other hand increased the PA amplitude compared to low value cues (mean ± s.e.m: 0.02 ± 0.2 and -0.86 ± 0.33, for the PA amplitude of high-compared to low-value cues, t(35) = 2.278, p = 0.029, d_z_ = 0.380).

Analysis of P1 component revealed only a main effect of modality (F_(1,35)_ = 5.33, P = 0.027, η_p_^2^ =0.132) but no main or interaction effect with reward value (both ps>0.1).

In N1 (170-250 ms) window, we found an interaction effect between reward value and modality (F(1,35) = 6.46, P = 0.016, η_p_^2^ = 0.156). Cross-modal high-value cues decreased the N1 negativity compared to low value cues (mean ± s.e.m: 1.21 ± 0.37 and -0.46 ± 0.41, t(35) = 3.245, p = 0.003, d_z_ = 0.541). The modulation of N1 amplitude was not significant for intra-modal cues.

In P3 window (300-600 ms), we found no main effect of modality, reward value or an interaction between reward and modality.

The analysis of the latency of P1 and N1 components revealed only a significant value × modality interaction effect for N1 latency (F(1,35) = 6.38, P = 0.016, η_p_^2^ = 0.154). Intra-modal, high value cues elicited faster N1 responses compared to low values cues (mean ± s.e.m: -4.94 ± 4.61 and 8.33 ± 3.39, for N1 latency of high- and low-value cues respectively corrected for pre-conditioning differences, t(35) = -2.228, p = 0.032, d_z_ = 0.371).

## Supplementary Figures

**Supplementary Figure 1.**
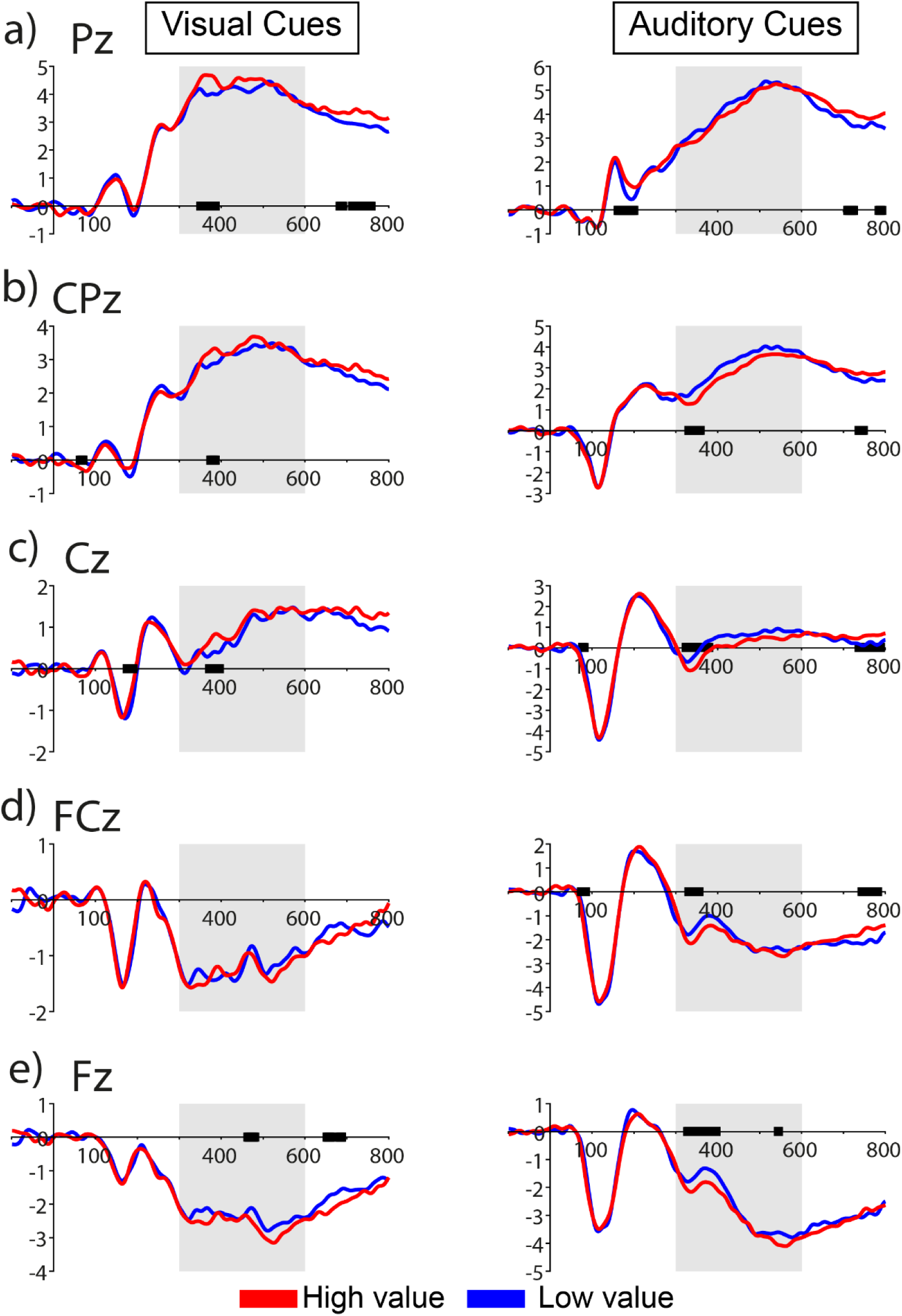
ERPs of midline electrodes during the reward associative learning (conditioning phase). Black marks on x axis correspond to times points when a significant difference between high and low reward value cues were found (uncorrected p<0.05), as measured by a moving window of 10 ms.

**Supplementary Figure 2.**
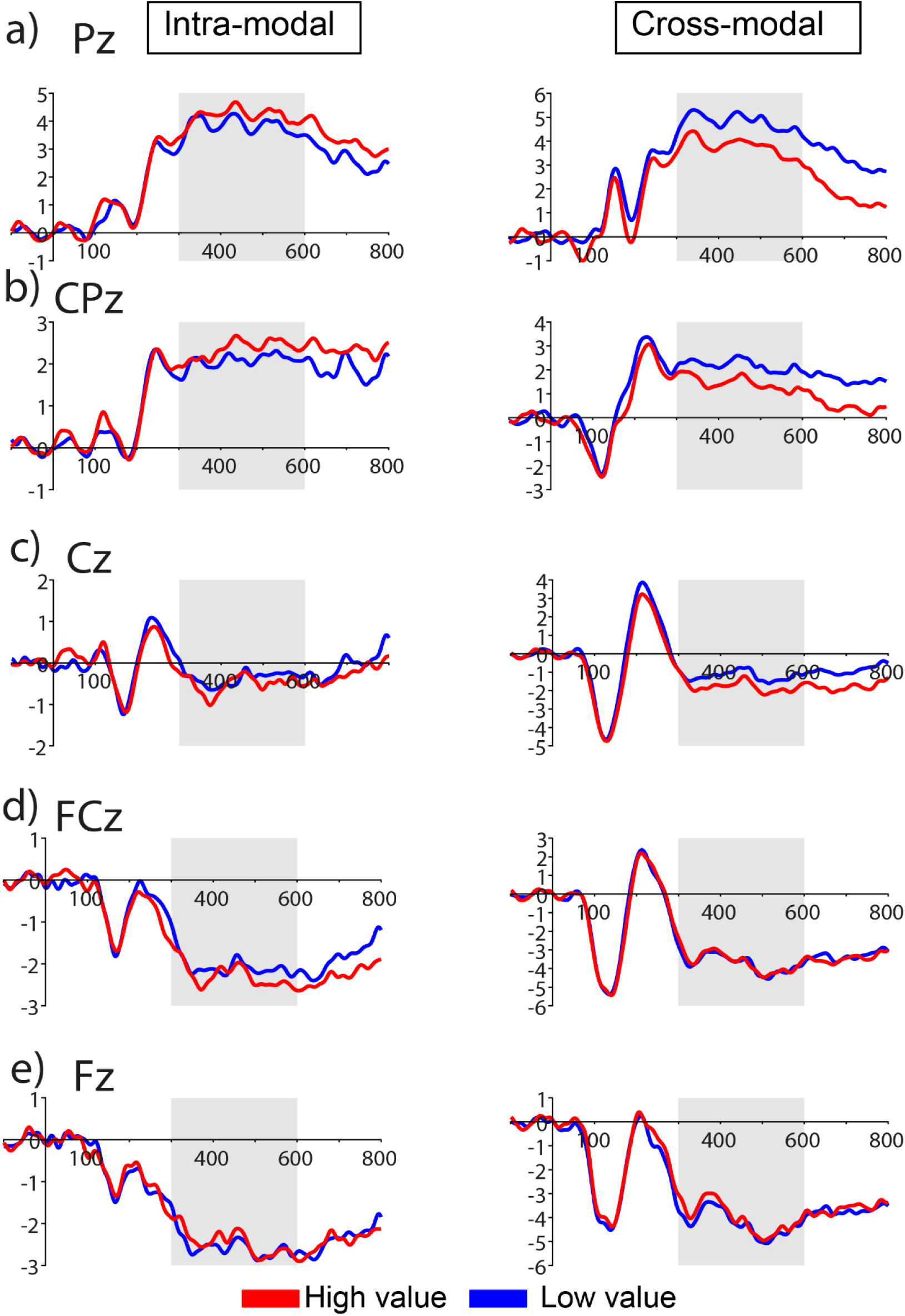
ERPs of midline electrodes during the pre-conditioning phase.

**Supplementary Figure 3.**
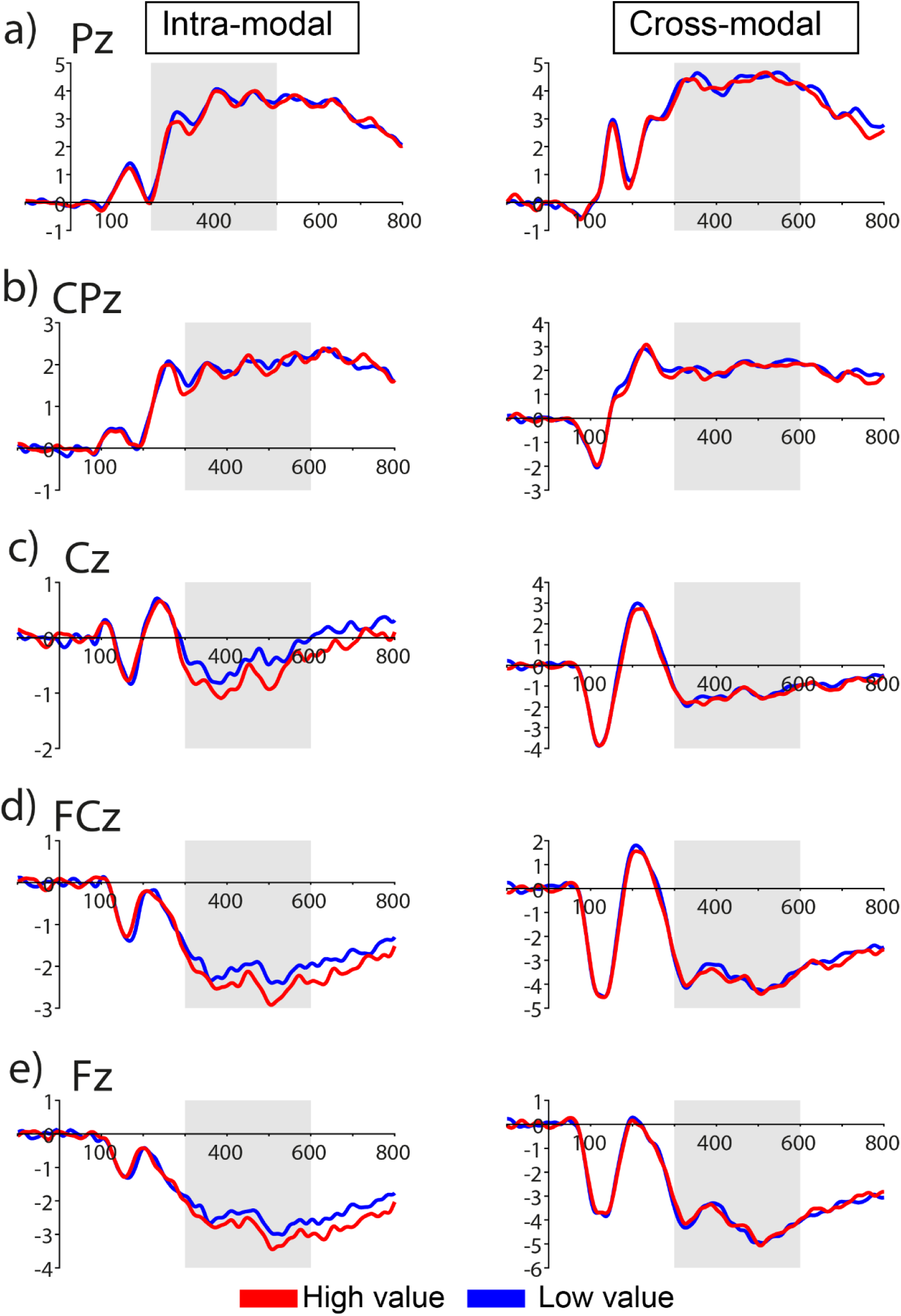
ERPs of midline electrodes during the post-conditioning phase.

**Supplementary Figure 4.**
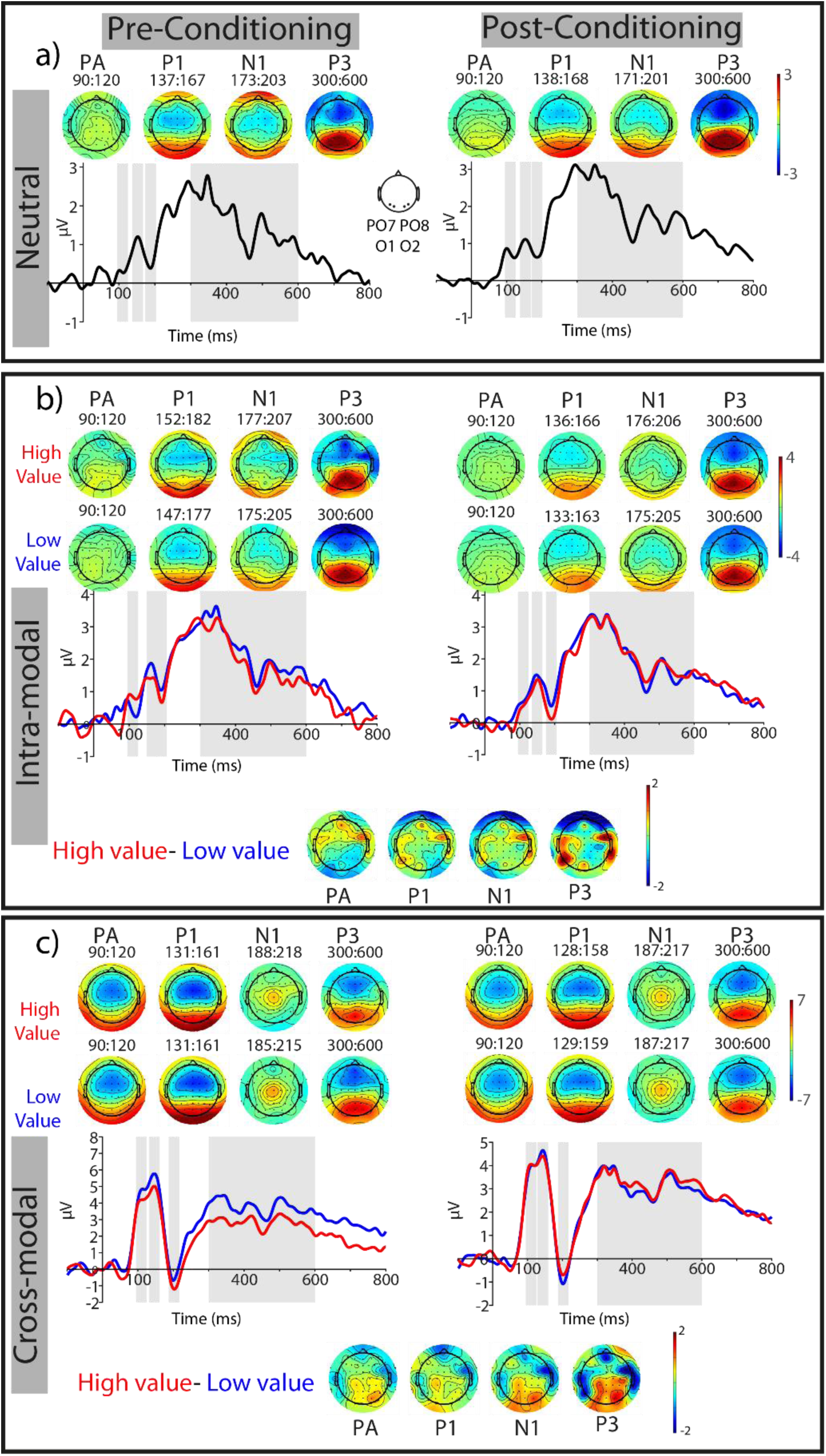
Contralateral responses of the posterior ROI during pre- and post-conditioing phases, see also Figure 4 in the main text.

**Supplementary Figure 5.**
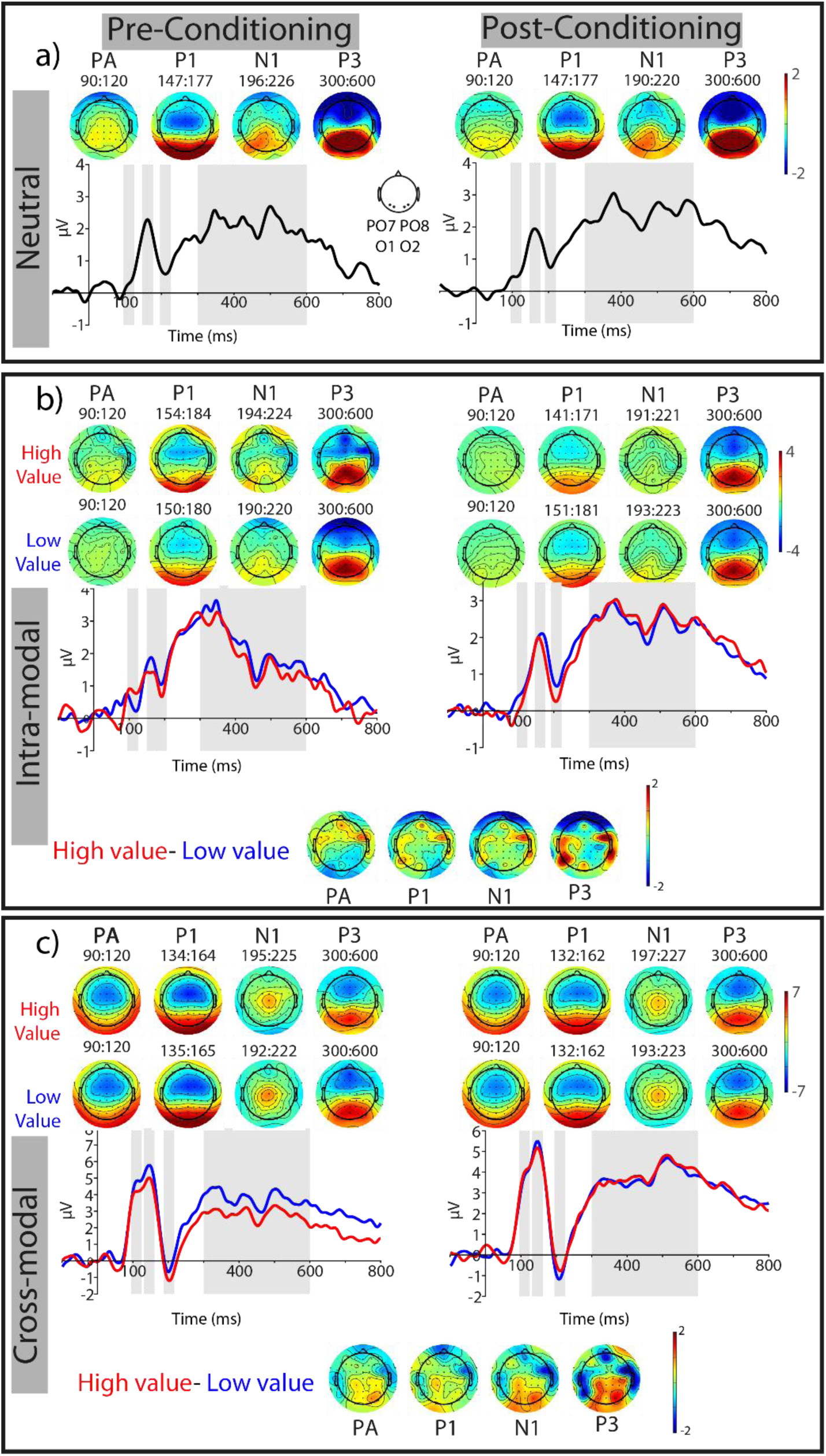
ERP results of the posterior ROI when only correct trials were included, see also Figure 4 in the main text.

